# The Alzheimer’s gene *SORL1* is a regulator of endosomal traffic and recycling in human neurons

**DOI:** 10.1101/2021.07.26.453861

**Authors:** Swati Mishra, Allison Knupp, Marcell P. Szabo, Charles A. Williams, Chizuru Kinoshita, Dale W. Hailey, Yuliang Wang, Jessica E. Young

**Author notes:** Equal Contribution (Swati Mishra, Allison Knupp).

## Abstract

**Background:** Loss of the Sortilin-related receptor 1 (*SORL1*) gene seems to act as a causal event for Alzheimer’s disease (AD). Recent studies have established that loss of *SORL1*, as well as mutations in autosomal dominant AD genes *APP* and *PSEN1/2*, pathogenically converge by swelling early endosomes, AD’s cytopathological hallmark. Acting together with the retromer trafficking complex, *SORL1* has been shown to regulate the recycling of the amyloid precursor protein (APP) out of the endosome, contributing to endosomal swelling and to APP misprocessing. We hypothesized that *SORL1* plays a broader role in neuronal endosomal recycling and used human induced pluripotent stem cell derived neurons (hiPSC-Ns) to test this hypothesis. We examined endosomal recycling of three transmembrane proteins linked to AD pathophysiology: APP, the BDNF receptor Tropomyosin-related kinase B (TRKB), and the glutamate receptor subunit AMPA1 (GLUA1).

**Methods:** We used isogenic hiPSCs engineered to have *SORL1* depleted or to have enhanced *SORL1* expression. We differentiated neurons from these cell lines and mapped the trafficking of APP, TRKB and GLUA1 within the endosomal network using confocal microscopy. We also performed cell surface recycling and lysosomal degradation assays to assess the functionality of the endosomal network in both *SORL1* depleted and overexpressing neurons. Finally, we analyzed alterations in gene expression in *SORL1* depleted neurons using RNA-sequencing.

**Results:** We find that as with APP, endosomal trafficking of GLUA1 and TRKB is impaired by loss of *SORL1*. We show that trafficking of all three cargo to late endosomes and lysosomes is affected by manipulating SORL1 expression. We also show that depletion of *SORL1* significantly impacts the endosomal recycling pathway for APP and GLUA1 at the level of the recycling endosome and trafficking to the cell surface. This has a functional effect on neuronal activity as shown by multi-electrode array (MEA). Conversely, increased *SORL1* expression enhances endosomal recycling for APP and GLUA1. Our unbiased transcriptomic data further support *SORL1*’s role in endosomal recycling. We observe altered expression networks that regulate cell surface trafficking and neurotrophic signaling in SORL1 depleted neurons.

**Conclusion:** Collectively, and together with other recent observations, these findings suggest that *SORL1* is a broad regulator of retromer-dependent endosomal recycling in neurons, a conclusion that has both pathogenic and therapeutic implications for Alzheimer’s disease.

## BACKGROUND

Alzheimer’s disease (AD) is a progressive neurodegenerative disorder and the most common cause of dementia. The underlying contributors to AD pathology encompass several biological pathways, including endosomal function, amyloid precursor protein (APP) processing, immune function, synaptic function, and lipid metabolism(Karch and Goate, 2015). Among these, endosomal dysfunction in neurons is emerging as a potential causal mechanism(Small and Petsko, 2020). Mutations in the amyloid precursor protein (*APP*) and the two presenilins (*PSEN1* and *PSEN2*) lead to early-onset autosomal dominant AD. When these mutations are modelled in human neurons and other systems they cause endosomal swelling, indicative of traffic jams, a phenotype that is a cytopathological hallmark of AD(Cataldo et al., 2000; Choi et al., 2013; Kwart et al., 2019). Recent genetic studies have identified a fourth gene, the trafficking receptor ‘sortilin related receptor 1’ (*SORL1*), which, when harboring frame-shift mutations leading to premature stop codons, is described as causal for AD(Holstege et al., 2017; Raghavan et al., 2018; Scheltens et al., 2021). Interestingly, *SORL1* is also linked to the more common, late-onset form of AD(Lambert et al., 2013; Rogaeva et al., 2007b) and its expression is lost in sporadic AD brains (Dodson et al., 2006; Thonberg et al., 2017). When modelled in human neurons, *SORL1* depletion phenocopies *APP* and *PSEN* mutations by causing endosomal swelling(Hung et al., 2021; Knupp et al., 2020).

The *SORL1* gene codes for the protein SORLA, an endosomal sorting protein that is also an adaptor molecule for the retromer trafficking complex (Fjorback et al., 2012; Rogaeva et al., 2007a; Small and Gandy, 2006). Retromer recycles cargo out of the early endosome, either from the endosome to the trans-Golgi network or, with greater importance for neurons, back to the cell surface(Fjorback et al., 2012; Seaman, 2012). To date, the best evidence for *SORL1*’s role in retromer-dependent endosomal recycling comes from studies investigating APP trafficking(Fjorback et al., 2012; Schmidt et al., 2007; Willnow and Andersen, 2013). Our previous work demonstrated that *SORL1* depletion retains APP in early endosomes, which may contribute to endosomal swelling by blocking recycling(Knupp et al., 2020).

Retromer-dependent trafficking in neurons, however, also recycles cargo other than APP. For example, retromer is required for the normal recycling of glutamate receptors, a trafficking event that mediates synaptic plasticity and synaptic health, and this dependency occurs independent of retromer’s role in APP recycling(Park et al., 2004; Temkin et al., 2017). Neurotrophin receptors are also trafficked through the endosomal system, in a retromer-dependent manner, and are important for synaptic health(Klinger et al., 2015; Patapoutian and Reichardt, 2001; Rohe et al., 2013).

Here we used human induced pluripotent stem cell derived-neurons (hiPSC-Ns) to test the hypothesis that *SORL1* plays a broader role in neuronal endosomal recycling. We use our previously described *SORL1*-depleted hiPSC lines to generate hiPSC-Ns, which model the loss of *SORL1* expression that occurs in AD(Knupp et al., 2020). Furthermore, we used previously established cell lines engineered to overexpress *SORL1* 2-3-fold over wild-type levels(Young et al., 2015) to test the effects of enhanced *SORL1* expression in hiPSC-Ns on these trafficking pathways. Importantly, all cell lines are isogenic. We map the trafficking effects these manipulations have on three specific receptors, APP, the GLUA1 subunit of the AMPA receptor, and neurotrophin receptor TRKB, all of which are implicated in AD (Devi and Ohno, 2015; Dewar et al., 1991; Ginsberg et al., 2019; Martin-Belmonte et al., 2020; Wakabayashi et al., 1999; Yasuda et al., 1995).

Finally, we performed RNA-sequencing on the *SORL1* depleted cell lines to explore an unbiased transcriptomics analysis induced by *SORL1* depletion. The results generally confirmed our hypothesis, showing that *SORL1* is a broad regulator of endosomal recycling in neurons, a conclusion that has both pathogenic and therapeutic implications.

## METHODS

### Cell lines

#### Cell lines generated by CRIPSR/Cas9 gene editing technology

The generation of the cell lines used in this paper is described in our previously published work (Knupp et al., 2020) and consists of four clones: Two wild-type clones, designated clone A6 and clone A7, and two *SORL1*KO clones, designated clone E1 and clone E4. Cell lines were generated from our previously published and characterized CV background human induced pluripotent stem cell line(Young et al., 2015). This cell line is male and has a APOE ε3/ε4 genotype(Levy et al., 2007). All four clones were shown to have normal karyotypes and are routinely tested for mycoplasma (MycoAlert). The clones used in the experiments in this work are listed in the figure legends.

#### CRISPR/Cas9 gRNA, ssODN, and Primer Sequences

gRNA: ATTGAACGACATGAACCCTC ssODN: GGGAATTGATCCCTATGACAAACCAAATACCATCTACATTGAACGACATGAACCCTCTGGC TACTCCACGTCTTCCGA AGTACAGATTTCTTCCAGTCCCGGGAAAACCAGGAAG Forward primer: ctctatcctgagtcaaggagtaac Reverse primer: ccttccaattcctgtgtatgc PCR amplifies 458 bp sequence. These sequences have been previously published in (Knupp et al., 2020)

#### SORL1 overexpression cell lines

Isogenic cell lines with overexpression of SORL1 were generated as previously described(Young et al., 2015). These lines are generated from the CV parental line, the same parental line as the *SORL1*KO cell lines were made from. Briefly, stable integration of SORL1 cDNA into the genome was achieved by using piggybac transposon system (Systems Biosciences). Vector alone (WT) or vector with SORL1 cDNA (*SORL1*OE) constructs were introduced into iPSCs by electroporation and stable cell lines were selected with puromycin (2ug/ml) treatment. We obtained one SORL1OE stable cell line and one vector alone stable cell line. For all overexpression experiments *SORL1*OE cells were compared to the vector alone controls.

### hiPSC Neuronal Differentiation

hiPSCs were differentiated to neurons using dual-SMAD inhibition(Chambers et al., 2009; Shi et al., 2012). Briefly, hiPSCs were plated on Matrigel coated 6-well plates at a density of 3.5 million cells per well and fed with Basal Neural Maintenance Media (1:1 DMEM/F12 + glutamine media/neurobasal media, 0.5% N2 supplement, 1% B27 supplement, 0.5% GlutaMax, 0.5% insulin-transferrin-selenium, 0.5% NEAA, 0.2% β-mercaptoethanol; Gibco, Waltham, MA) + 10mM SB-431542 + 0.5mM LDN-193189 (Biogems, Westlake Village, CA). Cells were fed daily for seven days. On day eight, cells were incubated with Versene, gently dissociated using cell scrapers, and passaged at a ratio of 1:3. On day nine, media was switched to Basal Neural Maintenance Media and fed daily. On day 13, media was switched to Basal Neural Maintenance Media with 20 ng/mL FGF (R&D Systems, Minneapolis, MN) and fed daily. On day sixteen, cells were passaged again at a ratio of 1:3. Cells were fed until approximately day twenty-three. At this time, cells were FACS sorted to obtain the CD184/CD24 positive, CD44/CD271 negative neural precursor cell (NPC) population. Following sorting, NPCs were expanded for neural differentiation. For cortical neuronal differentiation, NPCs were plated out in 10cm cell culture dishes at a density of 6 million cells/10cm plate. After 24 hours, cells were switched to Neural Differentiation media (DMEM-F12 + glutamine, 0.5% N2 supplement, 1% B27 supplement, 0.5% GlutaMax) + 0.02ug/mL brain-derived neurotrophic factor (PeproTech, Rocky Hill, NJ) + 0.02ug/mL glial-cell-derived neurotrophic factor (PeproTech) + 0.5mM dbcAMP (Sigma Aldrich, St Louis, MO). Media was refreshed twice a week for three weeks. After three weeks, neurons were selected for CD184/CD44/CD271 negative population by MACS sorting and plated for experiments. The data presented in this study represent 2-3 neuronal differentiations.

### Purification of Neurons

Following three weeks of differentiation, neurons were dissociated with accutase and resuspended in Magnet Activated Cell Sorting (MACS) buffer (PBS + 0.5% bovine serum albumin [Sigma Aldrich, St Louis, MO] + 2mM ethylenediaminetetraacetic acid [Thermo Fisher Scientific, Waltham, MA]). Following a modification of (Yuan et al., 2011), cells were incubated with PE-conjugated mouse anti-Human CD44 and mouse anti-Human CD184 antibodies (BD Biosciences, San Jose, CA) at a concentration of 5µl/10 million cells. Following antibody incubation, cells were washed with MACS buffer and incubated with anti-PE magnetic beads (BD Biosciences, San Jose, CA) at a concentration of 25µl/10 million cells. Bead-antibody complexes were pulled down using a rare earth magnet, supernatants were selected, washed, and plated at an appropriate density.

### DQ Red BSA assay for visualization of lysosomal degradation

Lysosomal proteolytic degradation was evaluated using DQ Red BSA (#D-12051; Thermo Fisher Scientific), a fluorogenic substrate for lysosomal proteases, that generates fluorescence only when enzymatically cleaved in intracellular lysosomal compartments. hiPSC-derived neurons were seeded at a density of 400,000 cells/well of a matrigel coated 48 well plate. After 24 hours, cells were washed once with DPBS, treated with complete media containing either 10µg/ml DQ Red BSA or vehicle (PBS) and incubated for 6 hours or 24 hours (Davis et al., 2021; Romano et al., 2021; Tian et al., 2015)at 37°C in a 5% CO_2_ incubator as described in(Marwaha and Sharma, 2017). At the end of 6 or 24 hours, cells were washed with PBS, fixed with 4% PFA and immunocytochemistry was performed as described in methods. Cells were imaged using a Leica SP8 confocal microscope and all image processing was completed with ImageJ software. Cell bodies were identified by MAP2 labeling, and fluorescence intensity of DQ Red BSA was measured in regions of the images containing the MAP2 label.

### Immunocytochemistry

For immunocytochemistry, cells were fixed with 4% PFA for 20 minutes. Fixed cells were washed three times with PBST (PBS with 0.05% tween 20), permeabilized with Triton X-100 in PBS for 15 minutes, washed twice again with PBST, blocked with 5% BSA in PBS at room temperature for 1h and incubated with appropriate primary antibodies overnight at 4°C. The next day, cells were incubated with appropriate secondary antibodies and 1µg/ml DAPI for 1 hour at RT, washed three times with PBST and mounted on glass slides with Prolong Gold Antifade mountant (#P36930; Thermo Fisher Scientific).

### Colocalization analysis

To investigate colocalization with endo-lysosomal compartments, hiPSC-derived neurons were labeled with markers specific for each intra-cellular compartment (EEA1 for early endosomes, Rab7 for late endosomes, LAMP1 for lysosomes and Rab11 for recycling endosomes) using immunocytochemistry. A minimum of 10 fields of confocal z-stack images were captured under blinded conditions using a Yokogawa W1 spinning disk confocal microscope (Nikon) and a 100X plan apochromat oil immersion objective. Median filtering was used to remove noise from images and Otsu thresholding was applied to all images. Colocalization was quantified using the JACOP plugin(Bolte and Cordelieres, 2006) in Image J software(Schindelin et al., 2012) and presented as Mander’s correlation coefficient.(Dunn et al., 2011; Manders et al., 1993). For all imaging experiments the data was analyzed in a blinded manner.

### Cell Surface Staining

Cell surface expression of GLUA1 and APP was determined using immunocytochemistry and confocal microscopy. To label proteins at the cell surface, cells were fixed with 4% PFA, washed and treated with primary and secondary antibodies as described in the ‘immunocytochemistry’ section of methods. Permeabilization with 0.1% Triton X-100 was not performed for this experiment. To label total protein levels, cells were fixed with 4% PFA, washed, permeabilized with 0.1% Triton X-100 and treated with primary and secondary antibodies as described in the ‘immunocytochemistry’ section of methods. Analysis of fluorescence intensity was done using Image J software. Cell surface expression was represented as ratio of fluorescence intensity measured under non-permeabilized conditions and fluorescence intensity measured under permeabilized conditions. For all imaging experiments the data was analyzed in a blinded manner.

### Multielectrode (MEA) assay

hiPSC-derived neural progenitor cells were differentiated into neurons and neurons were purified as previously described in methods. Purified neurons were mixed with unpurified neurons in a ratio of 5:1 and this mixture was plated onto a matrigel coated 48 well MEA plate (Axion Biosystems; # M768-tMEA-48W) at a cell density of 8000 cells/ul (total number of cells/well = 50,000). MEA-plated neurons were initially cultured in neural differentiation media. Media was gradually switched to BrainPhys media (Stem cell technologies; # 05790) by replacing half of a well’s media twice a week. BrainPhys media was supplemented with B27, N2, BDNF, GDNF, and db-cAMP.

### Multielectrode (MEA) analysis

Electrical signals from neurons in the MEA plates were recorded twice a week using Axion Biosystems Maestro Pro system. Signals were recorded at a sampling frequency of 12.5 kHz with a 3 kHz Kaiser Window low pass filter and 200 Hz high pass filter. Spikes were detected using Axion Axis Navigator recording software with the adaptive threshold method. Recordings were analyzed using the Axion Neural Metric Tool. Firing rate data were limited to active electrodes that detected a minimum of five spikes a minute. Firing rate data from all active electrodes in a well were averaged for plotting and statistical testing.

### Antibodies

The following primary antibodies were used: Early endosome antigen 1 (EEA1) at 1:500 (#610456; BD Biosciences); amyloid precursor protein (APP) at 1:500 (#ab32136; Abcam); microtubule-associated protein 2 (MAP2) at 1:1000 (ab92434; Abcam); Ras-related protein Rab-7a (Rab7) at 1:1000 (ab50533; Abcam); Ras-related protein Rab-11 (Rab11) at 1:250 (#610656; BD Biosciences); Lysosome associated membrane protein-1 (LAMP1) at 1:250 (#sc 2011; Santa Cruz); Tropomyosin receptor kinase B (TRKB; # ab18987; abcam) at 1:1000, GLUA1(# MAB2263; Millipore sigma) at 1:500 andVPS35(Abcam; #ab97545) at 1:500. DAPI was used at a final concentration of 0.1µg/ml (Alfa Aesar).

### Transferrin recycling assay

To measure recycling pathway function, we utilized transferrin recycling assay as previously described(Rapaport et al., 2010). Purified neurons were seeded at 400,000 cells/well of a 24 well plate containing matrigel coated 12 mm glass coverslip/well. After 5 DIV, cells were washed once with DMEM-F12 medium and incubated with starving medium (DMEM-F12 medium+25mM HEPES+ 0.5% BSA) for 30 minutes at 37°C in a 5% CO_2_ incubator to remove any residual transferrin. Thereafter, cells were pulsed with either 100µg/ml transferrin from human serum conjugated with Alexa Fluor™ 647(#T23366; Thermo Fisher Scientific) or vehicle (PBS) in ‘starving medium’. At the end of 10 mins, cells were washed twice with ice-cold PBS to remove any external transferrin and stop internalization of transferrin and washed once with acid stripping buffer (25mM citric acid+24.5mM sodium citrate+280mM sucrose+0.01mM Deferoxamine) to remove any membrane bound transferrin. Next, cells were either fixed in 4% PFA or ‘Chase medium’ (DMEM-F12+50µM Deferoxamine+20mM HEPES+500µg/ml Holo-transferrin) was added for different time points. Immunocytochemistry was done using MAP2 antibody to label neurons, confocal images were captured using Leica SP8 confocal microscope under blinded conditions. Fluorescence intensity of transferrin was measured using ImageJ software. Recycling function was presented as transferrin fluorescence intensity as a percentage of the fluorescence intensity measured at time zero.

### Measurement of lysosome and recycling endosome size

Immunocytochemistry using antibodies for LAMP1 and MAP2 or RAB11 and MAP2 was performed as described above. Using a Leica SP8 confocal microscope with an apochromatic 63X oil immersion lens, z-stack images were obtained under blinded conditions. For the LAMP1 analysis, 17-34 fields were analyzed for a total of **45 – 76** cells analyzed. For the RAB11 analysis, 15-30 fields were analyzed for a total of **59 – 124** cells analyzed. Vesicle size measurements were performed using Cell Profiler software as previously described (Knupp et al., 2020) (McQuin et al., 2018). Briefly, the vesicle channel was masked using the MAP2 channel and automated segmentation algorithms were used to identify individual puncta. The pixel area of each puncta was measured and is presented as mean area of all puncta per field.

### Statistical Analysis

For all imaging experiments, data was collected and analyzed in a blinded manner. Data was assessed for significance using parametric two-tailed unpaired Student’s t-tests or two-way ANOVA tests. Data is represented as mean ± standard deviation to show the spread of the data.

Significance was defined as a value of p ≤ 0.05. All statistical analysis was completed using GraphPad Prism software. Statistical details of individual experiments, including biological and technical replicate information, can be found in the figure legends. All of the raw statistical data for the experiments in this paper, including means, difference between the means ± SEM, and 95% confidence intervals are presented in Supplemental Table 1.

### RNA-sequencing analysis

#### RNA Extraction

RNA was collected from 3 separate differentiations including a combination of two WT clones and two *SORL1*KO clones. Each sample includes 2-3 technical replicates. RNA was collected from 2 million purified neurons for each sample. Purification of total RNA was completed using the PureLink RNA Mini Kit (Thermo Fisher 12183018A). Assessment of purified RNA was completed using a NanoDrop. Final RNA quantification was completed using the Quant-iT RNA assay (Invitrogen) and RNA integrity analysis was completed using a fragment analyzer (Advanced Analytical).

#### Library Prep and Sequencing

Library preparation was completed using the TruSeq Stranded mRNA kit (Illumina RS-122-2103) per manufacturer instructions. Sequencing was performed on a NovaSeq 6000 instrument.

#### Data analysis

Raw reads were aligned to GRCh38 with reference transcriptome GENCODE release 29 using STAR v2.6.1d(Dobin et al., 2013). Gene-level expression quantification is generated by RSEM v1.3.1(Li and Dewey, 2011). Genes with fewer than 20 normalized reads across all samples were omitted from further analysis. We did observe variation in the transcriptome based on differentiation (Supplemental Figure 3A), however this was corrected for using the sva package(Leek et al., 2012).

To identify differentially expressed genes (DEGs), we used DESeq(Anders and Huber, 2010). Briefly, we fit two models: a null model where gene expression only depends on batch effects (i.e., differentiation), and an alternative model where gene expression depends on both genotype (*SORL1*KO vs. WT) and batch effects. Chi-squared tests were performed to compare both fits, and we declare a gene as differentially expressed only when the alternative model fits the expression data better. DEGs are defined as genes with false discovery rate less than 0.05 and fold change greater than 1.5. The top gene ontology package(Alexa et al., 2006) and the SynGO synaptic gene ontology annotations(Koopmans et al., 2019) were used to identify GO terms that were enriched. GO terms were tested according to the Fisher’s exact test. Finally, we mapped DEGs onto receptor-ligand interaction diagrams generated by Ramilowski et al(Ramilowski et al., 2015) using the igraph plugin(Csardi G, 2006). To compare the amount of down-regulated vs. up-regulated genes we used a 2-sample test for equality of proportions with continuity correction in R.

## RESULTS

### *SORL1* depletion increases neuronal cargo localization in early endosomes

Using CRISPR/Cas9 genome editing techniques, we previously generated hiPSC-derived neurons (hiPSC-Ns) deficient in *SORL1* expression due to indels introduced in exon 6. We demonstrated that loss of *SORL1* expression in these neurons leads to enlarged early endosomes and an increased colocalization of APP within early endosomes, indicative of endosomal traffic jams(Knupp et al., 2020). We utilized these same cell lines (hereafter referred to as *SORL1*KO and their guide-matched isogenic wild-type clones referred to as WT) to examine localization of the BDNF receptor TRKB and the GLUA1 subunit of the neuronal AMPA receptor. TRKB has been shown to bind to SORLA and this interaction mediates trafficking of TRKB to synaptic plasma membranes(Rohe et al., 2013). GLUA1 is trafficked via the retromer complex, of which SORLA is an adaptor protein(Fjorback et al., 2012; Temkin et al., 2017) and both of these cargo are important in maintaining healthy neuronal function. Because we previously observed an increase in APP localization in early endosomes, resulting in a decrease in localization in downstream vesicles such as Ras-related protein (Rab)7+ late endosomes with *SORL1* depletion(Knupp et al., 2020), we performed an immunocytochemical analysis of both TRKB and GLUA1 localization with the early endosome marker EEA1. Similar to our previous observations for APP, we documented significantly increased localization of both TRKB (Figure 1A) and GLUA1 (Figure 1B) in early endosomes in *SORL1*KOneurons as compared to isogenic WT control neurons. Accumulation of neuronal cargo in early endosomes is indicative of endosomal traffic jams, which are thought to impact the transit of cellular cargo through other arms of the endo-lysosomal network. Due to SORLA’s role as an adaptor protein for the retromer complex, we also examined whether *SORL1* depletion led to changes in retromer subunit localization. We observed that VPS35, a core subunit of the retromer cargo recognition complex is also mis-localized to early endosomes, similar to what we observed for APP, TRKB, and GLUR1 (Supplemental Figure 1).

**Figure 1.**
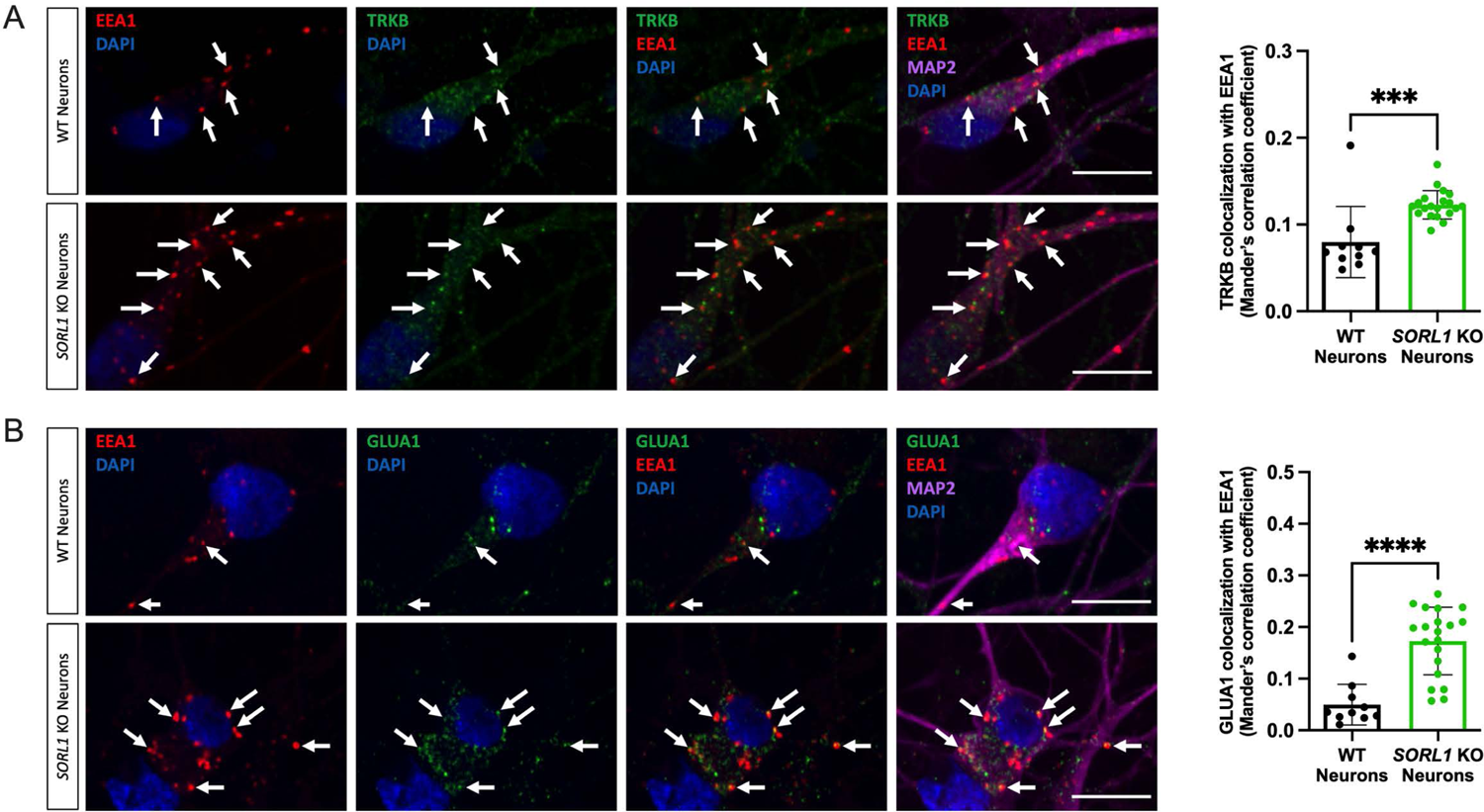
Loss of *SORL1* expression leads to increased TRKB and GLUA1 localization in early endosomes. Representative immunofluorescent images of WT and *SORL1*KO neurons showing increased colocalization of **(a)** TRKB (green) and **(b)** GLUA1 (green) with EEA1 (red). All neurons were immunolabeled with MAP2 (far-red) and counterstained with DAPI (blue). Scale bar: 10μm. In all cases, quantification of colocalization was represented as Mander’s correlation co-efficient (MCC). 1 WT and 2 *SORL1*KO isogenic clones were used for these experiments and 10 images per clone per genotype were analyzed. Data represented as mean ± SD. Data was analyzed using parametric two-tailed unpaired t test. Significance was defined as a value of *p < 0.05, **p < 0.01, ***p < 0.001, and ****p < 0.0001.

### Modulating *SORL1* expression impacts cargo trafficking throughout the endo-lysosomal system

The early endosome serves as a hub in which internalized cargo can be retrogradely transported to the trans-Golgi, recycled back to the cell surface or degraded as endosomes mature into late endosomes and lysosomes(Mayle et al., 2012). We have previously observed that APP localization within the trans-Golgi network was decreased in *SORL1*KO neurons(Knupp et al., 2020). Here we tested whether trafficking to the degradative arm of the endo-lysosomal network was affected in our *SORL1*KO neurons. Trafficking of substrates out of the early endosome to late endosomes and, subsequently, lysosomes is important for protein degradation and SORLA has been previously implicated in promoting Aβ degradation via lysosomes(Caglayan et al., 2014). We treated *SORL1* deficient neurons with DQ Red BSA, a proteolysis sensitive fluorogenic substrate that generates fluorescence only when enzymatically cleaved in intracellular lysosomal compartments. Since substrate degradation primarily occurs in lysosomes, altered fluorescence intensity of this reagent is a readout of altered lysosomal degradation(Marwaha and Sharma, 2017). We treated neurons with DQ Red BSA for 6 and 24 hours and analyzed fluorescence intensity using confocal microscopy. Consistent with loss of *SORL1* leading to endosomal traffic jams, we observed a significant reduction of DQ Red BSA fluorescence intensity at both time points in *SORL1*KO neurons compared to isogenic WT controls (Figure 2A). We next performed immunocytochemical staining to quantify the colocalization of our selected neuronal cargo with Rab7, a marker of late endosomes, and LAMP1 (Lysosomal Associated Membrane Protein 1), a lysosome marker. We show a significant decrease in co-localization of TRKB (Figure 2B) and GLUA1 (Figure 2C) with Rab7. This result is consistent with our previous observation for APP (Knupp et al., 2020). We analyzed colocalization of these cargo with LAMP1 and we observed a significant decrease with APP (Figure 2D) and TRKB (Figure 2E) and a trend of a decrease with GLUA1 (Figure 2F). These data indicate some fluidity in the network but suggest that trafficking of APP, TRKB and GLUA1 to late endosomes/lysosomes is all decreased by *SORL1*KO, although GLUA1 may be more likely to be trafficked to cell surface pathways or utilizes other adaptor proteins for late endosome to lysosomal trafficking. These changes in localization are not due to changes in expression of cargo. We have previously shown that APP levels do not change in *SORL1*KO neurons(Knupp et al., 2020) and also show here that protein expression of TRKB and GLUA1 are not different (Supplemental Figure 3).

**Figure 2.**
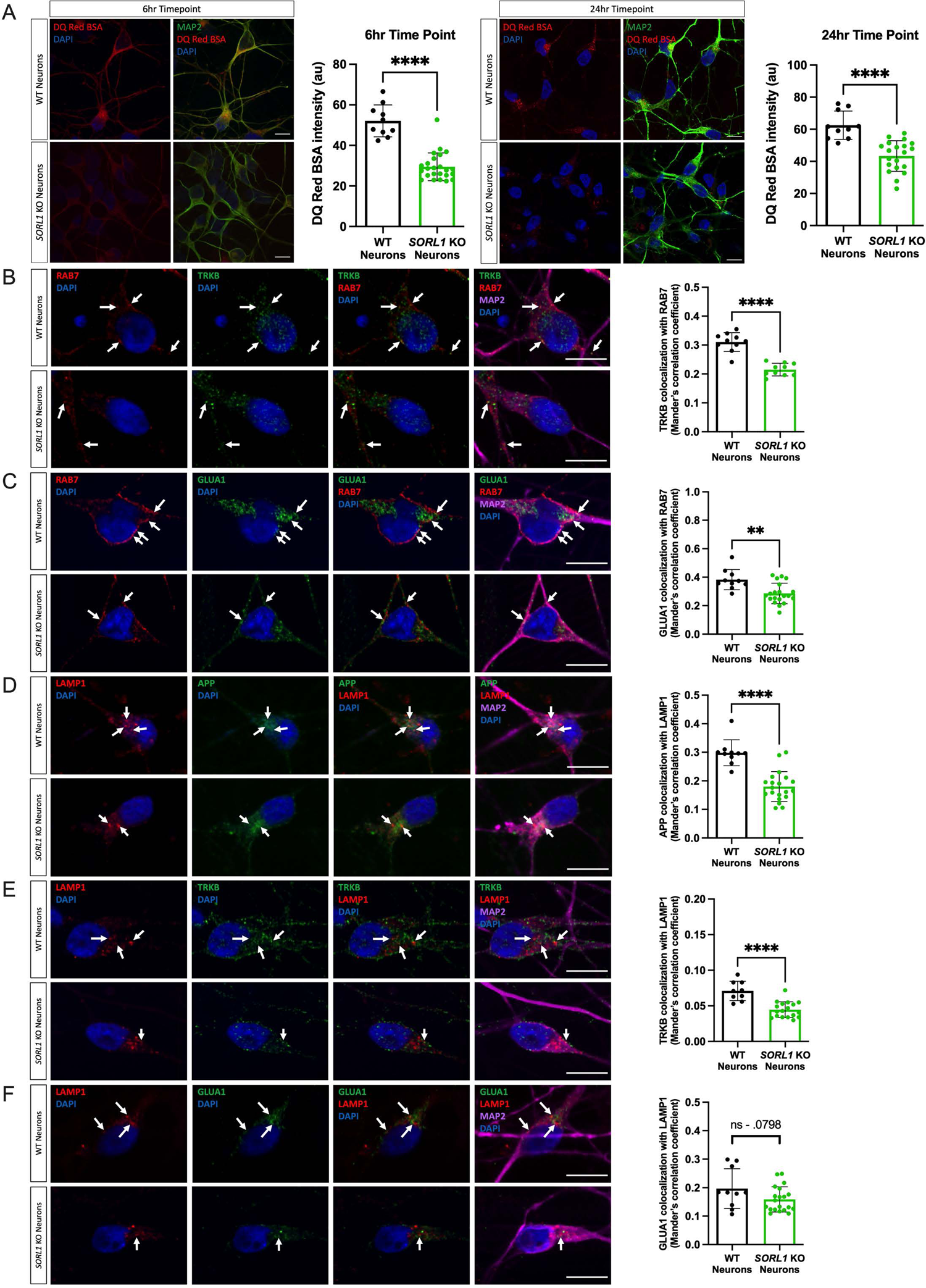
Loss of *SORL1* expression impairs trafficking to late endosomes and lysosomes. **a)** *SORL1*KO neurons show reduced lysosomal proteolytic activity as measured by DQ Red BSA. Representative immunofluorescent images of WT and *SORL1*KO neurons showing double immunolabeling for MAP2 (green) and DQ Red BSA (red). Scale bar: 10μm Quantification of fluorescence intensity of DQ Red BSA using ImageJ software. **(b-f)** *SORL1*KO neurons show reduced colocalization of cargo with late endosomes and lysosomes. Representative immunofluorescent images of WT and *SORL1*KO neurons showing reduced colocalization of **(b)** TRKB (green) and **(c)** GLUA1 (green) with Rab7 positive late endosomes (red) in *SORL1*KO neurons. Representative immunofluorescent images of WT and *SORL1*KO neurons showing reduced colocalization of **(d)** APP (green) and **(e)** TRKB (green) and **(f)** GLUA1 (green) with LAMP1 positive lysosomes (red) in *SORL1*KO neurons. Scale bar: 10μm. In all cases, quantification of colocalization was represented as Mander’s correlation co-efficient (MCC). 1 WT and 2 *SORL1*KO isogenic clones were used for these experiments and 10 images per clone per genotype were analyzed. Data represented as mean ± SD. Data was analyzed using parametric two-tailed unpaired t test. Significance was defined as a value of *p < 0.05, **p < 0.01, ***p < 0.001, and ****p < 0.0001.

We next utilized previously generated cell lines that overexpress *SORL1* cDNA using the piggybac transposon system(Young et al., 2015) to test whether increased *SORL1* expression may enhance the trafficking pathways that are impaired in the *SORL1*KO neurons. Importantly, the *SORL1* overexpressing (*SORL1*OE) cell line and control were generated in the same genetic background as our *SORL1*KO and isogenic WT cell lines. Interestingly, while there was no effect of *SORL1* overexpression on DQ Red BSA fluorescence at the earlier time point (6 hours), we did see a significant enhancement of DQ RED BSA trafficking at the 24-hour time point (Figure 3A). In accordance, we observed significantly increased localization of our studied cargo with late endosomal and lysosomal markers in the *SORL1*OE neurons (Figure 3 B-G).

**Figure 3:**
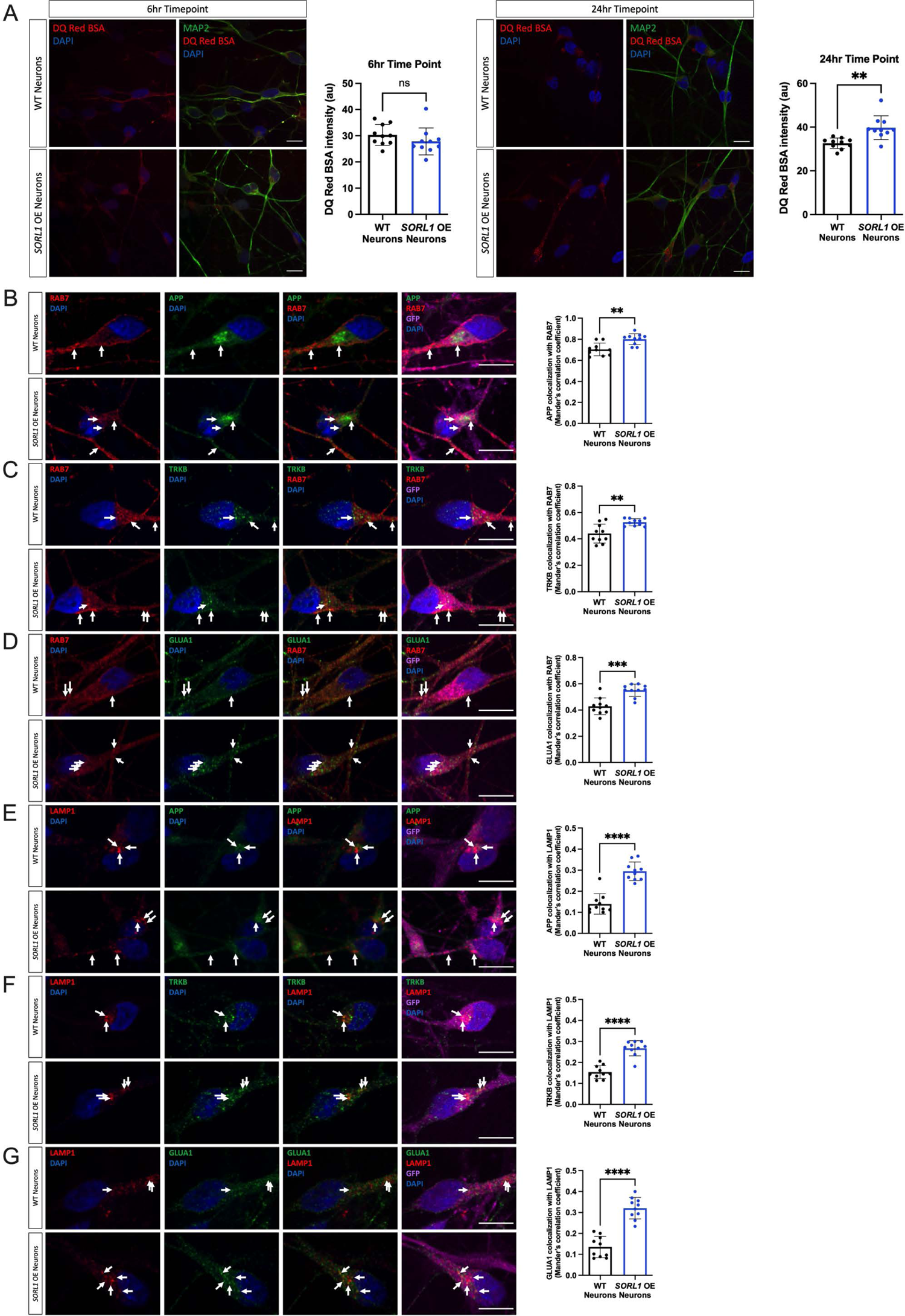
Enhancing *SORL1* Expression improves trafficking to late endosomes and lysosomes. **a)** *SORL1*OE neurons show no change in lysosomal proteolytic activity as measured using DQ Red BSA after a 6 hour treatment but do show an enhancement of trafficking at 24 hours. Representative immunofluorescent images of WT and *SORL1*OE neurons showing double immunolabeling for MAP2 (green) and DQ Red BSA (red). Scale bar: 10 μm. Quantification of fluorescence intensity of DQ Red BSA using ImageJ software. 1 cell line of each genotype (WT vs. SORL1 OE) were used for these experiments. 10 images per genotype were analyzed. Representative immunofluorescent images of WT and *SORL1*OE neurons showing increased colocalization of APP, TRKB and GLUA1 (green) with Rab7 (red) **(b-d)** and LAMP1 (red) **(e-g)** in *SORL1*OE neurons. *SORL1*OE neurons and controls have endogenous GFP expression due to the piggybac vector system. GFP fluorescence is pseudo-colored (Far-red) and was used to outline cell bodies. Scale bar: 10μm. Nuclei are counterstained with DAPI (blue). In all cases, quantification of colocalization was represented as Mander’s correlation co-efficient (MCC). 1 cell line of each genotype (WT vs. SORL1 OE) were used for these experiments. 10 images per genotype were analyzed. Data represented as mean ± SD. Data was analyzed using parametric two-tailed unpaired t test. Significance was defined as a value of *p < 0.05, **p < 0.01, ***p < 0.001, and ****p < 0.0001.

Lysosome size can influence lysosome function and is altered in AD(de Araujo et al., 2020; Hwang et al., 2019). Similarly, location and number of lysosomes within neurons can alter degradative activity (Cheng et al., 2018; Farfel-Becker et al., 2019; Gowrishankar et al., 2015; Yap et al., 2018) and in some cases, altered lysosomal distribution may represent an early neuropathological defect(Zigdon et al., 2017).Recently, loss of *SORL1* in hiPSC neurons was shown to contribute to lysosome dysfunction as indicated by both increased lysosome size and number as well as decreased cathepsin-D activity(Hung et al., 2021). Therefore, we first analyzed LAMP1-immunopositive puncta and also documented a significant increase in lysosome size and number in our *SORL1*KO neurons (Supplemental Figure 2A). Interestingly, although the number of lysosomes marked by LAMP1 puncta is increased in *SORL1KO* neurons, we did not observe a significant change in LAMP1 protein expression (Supplemental Figure 3). This may be partially explained by differences in autophagy in SORL1 KO neurons which we did not examine in this study but that has been previously reported(Hung et al., 2021) and further underscores the dynamic complexity of the endo-lysosomal network.

Next, we analyzed co-localization of Cathepsin-D and LAMP1 to determine if loss of *SORL1* expression leads to altered Cathepsin-D trafficking in neurons. Retromer trafficking is required to deliver one of the most abundant lysosomal proteases, pro-cathepsin D, to lysosomes via the mannose-6-phosphate receptor (M6PR)(Qureshi et al., 2018; Seaman, 2004). The SORLA protein has GGA domains similar to that of M6PR(Spoelgen et al., 2006), and mis-trafficking of Cathepsin-D to lysosomes could affect the maturation and degradative capacity of these organelles. Therefore, we analyzed co-localization of Cathepsin-D and LAMP1 to determine if loss of *SORL1* expression leads to altered Cathepsin-D trafficking in neurons. However, we did not observe a change in Cathepsin-D colocalization between WT and *SORL*1KO (Supplemental Figure 2B).

Taken together, our data suggest that *SORL1* loss in neurons reduces trafficking of cargo out of the early endosome to the late endosome and lysosome, contributing to lysosome stress as evidenced by an increase in size and number in these conditions while the decreased cathepsin-D activity observed upon *SORL1* loss(Hung et al., 2021) may not be due to impairment of lysosomal trafficking of the enzyme.

### Loss of *SORL1* impacts the endosomal recycling pathway

Another route out of the early endosome is via the endocytic recycling complex (ERC) which can send cargo either to the cell surface or to the trans-Golgi network(Grant and Donaldson, 2009; Mallard et al., 1998; Marsh et al., 1995; Maxfield and McGraw, 2004). To directly examine if *SORL1* expression alters recycling function, we performed a transferrin recycling assay using confocal microscopy. Transferrin can be recycled via a fast pathway within approximately 5-10 minutes after being internalized or via a slower pathway involving the ERC over longer periods of time(Ouellette and Carabeo, 2010; Sonnichsen et al., 2000). We examined the fluorescence intensity of Alexa Fluor 647-conjugated transferrin over a 40-minute time course in WT and *SORL1*KO neurons and observed that a higher percentage of intracellular fluorescent transferrin persisted in *SORL1*KO neurons at both early and later time points as compared to WT neurons, indicating reduced recycling pathway function (Figure 4A). Cargo destined for the cell surface can transit the ERC via Rab11+ recycling endosomes (Ren et al., 1998). Altered size of recycling endosomes can be indicative of dysfunctional recycling of cargo through these compartments. We tested whether loss of *SORL1* expression affected the size of Rab11+ recycling endosomes. Interestingly, we observed a significant increase in the size of Rab11+ recycling endosomes, although there was no change in Rab11 protein expression in the *SORL1*KO neurons (Figure 4B, Supplemental Figure 3), suggesting that this endosomal compartment is also under stress. To test if increased size is due to abnormal cargo trafficking through recycling endosomes, we assessed colocalization of APP, TRKB and GLUA1 with Rab11 and observed increased co-localization of all three cargo with Rab11+ structures in *SORL1*KO neurons compared to WT neurons (Figure 4C-E). Together, these data demonstrate that loss of *SORL1* impacts neuronal recycling endosome pathways by causing traffic jams in the recycling endosomes, similar to the effect that *SORL1* loss has on early endosomes.

**Figure 4.**
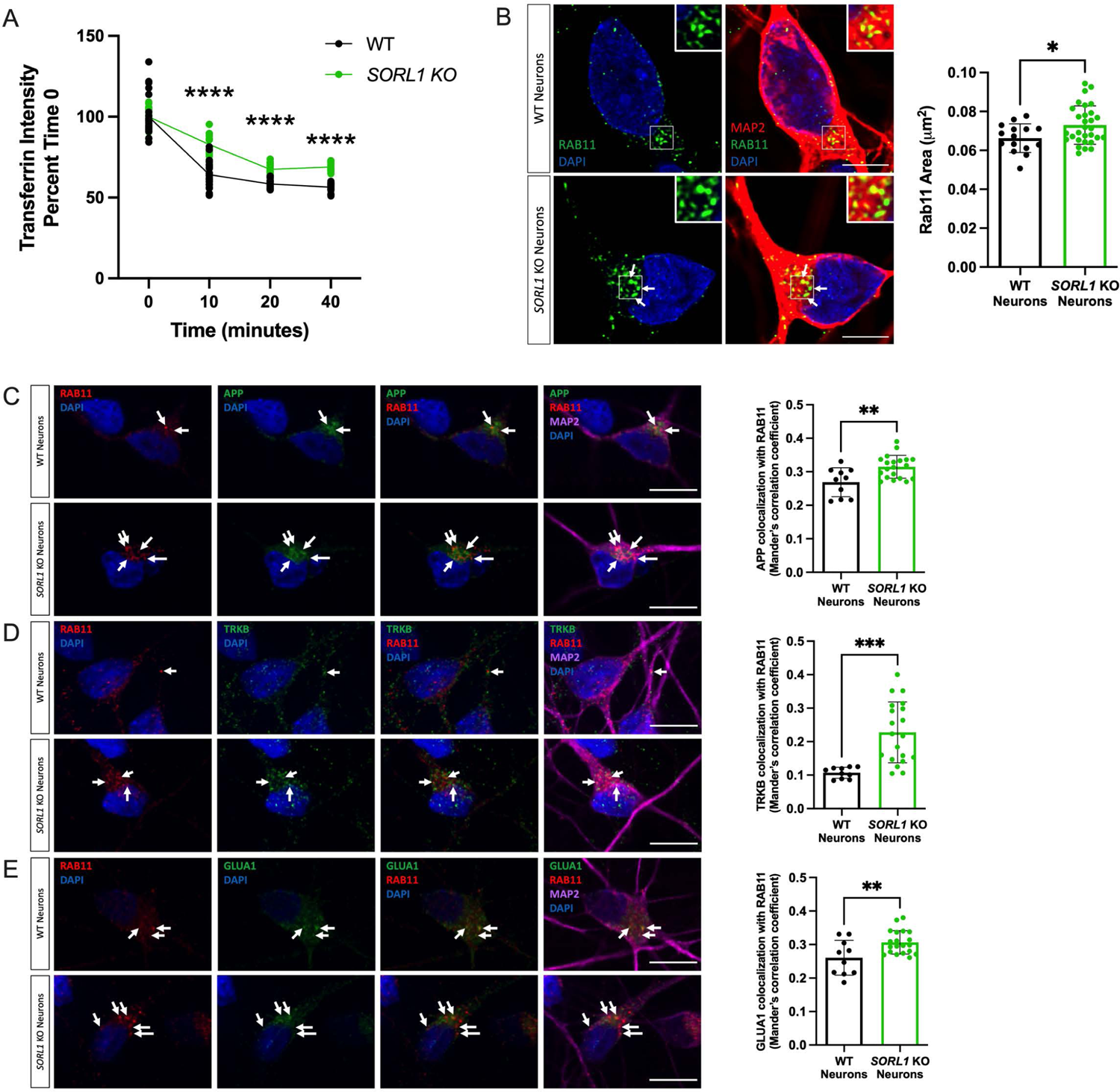
Loss of *SORL1* impacts the cell surface recycling pathway. **a)** *SORL1*KO neurons show slower rate of transferrin recycling. Quantification of fluorescence intensity of intracellular transferrin at different time points after treating cells with Alexa Fluor 647-conjugated transferrin for 15 mins, using ImageJ software. Data represented as percent of time 0 fluorescence intensity. 2 WT and 2 *SORL1*KO isogenic clones were used for these experiments. 12 images per clone per genotype were analyzed. **(b)** *SORL1*KO neurons show larger recycling endosomes. Representative immunofluorescent images of WT and *SORL1*KO neurons labeled with antibodies for MAP2 (red) and Rab11 (green). Nuclei were counterstained with DAPI (blue). Scale bar: 5μm. Quantification of size of Rab11 labeled recycling endosomes using CellProfiler software. 1 WT and 2 *SORL1*KO isogenic clones were used for these experiments. 15 images per clone per genotype were analyzed. Representative immunofluorescent images of WT and *SORL1*KO neurons showing increased colocalization of **(c)** APP (green), **(d)** TRKB (green) and **(e)** GLUA1 (green) with Rab11 positive recycling endosomes (red) in *SORL1*KO neurons. Scale bar:10μm In all cases, quantification of colocalization was represented as Mander’s correlation co-efficient (MCC). 1 WT and 2 *SORL1*KO isogenic clones were used for these experiments. 10 images per clone per genotype were analyzed. Data represented as mean ± SD. Data was analyzed using parametric two-tailed unpaired t test and two-was ANOVA. Significance was defined as a value of *p < 0.05, **p < 0.01, ***p < 0.001, and ****p < 0.0001.

### *SORL1* depletion reduces cell surface levels of cargo

Together, our data indicate that *SORL1*KO neurons have impaired cargo recycling with increased retention of cargo in recycling endosomes. These observations led us to test whether this cargo was indeed trafficked to the cell surface. A portion of APP has been shown to return to the cell surface via recycling endosomes(Das et al., 2016) and SORLA can interact with the sorting nexin SNX27 to return APP to the cell surface(Das et al., 2016; Huang et al., 2016), although in that study the exact compartment was not described. Furthermore, recycling endosomes are the source for AMPA receptors during long-term potentiation(Park et al., 2004). We therefore examined cell surface levels of APP and GLUA1 using immunofluorescence and confocal microscopy. We documented a significant decrease in cell surface staining of both APP (Figure 5A) and GLUA1 (Figure 5B) in *SORL1*KO neurons as compared to WT, consistent with our hypothesis that SORLA is involved in regulating traffic from recycling endosomes. Due to the importance of GLUA1 in the formation of functional excitatory synapses, we next analyzed neuronal activity by culturing *SORL1*KO and isogenic WT neurons on multi-electrode array (MEA) plates. Interestingly, we observed an early increase in the mean firing rate of *SORL1*KO neurons at an early time point (27 days post-plating) however neuronal firing in *SORL1*KO neurons was significantly reduced at a later time point (66 days post-plating) (Figure 5C) suggesting that synaptic activity may be partially impaired in these cells.

**Figure 5.**
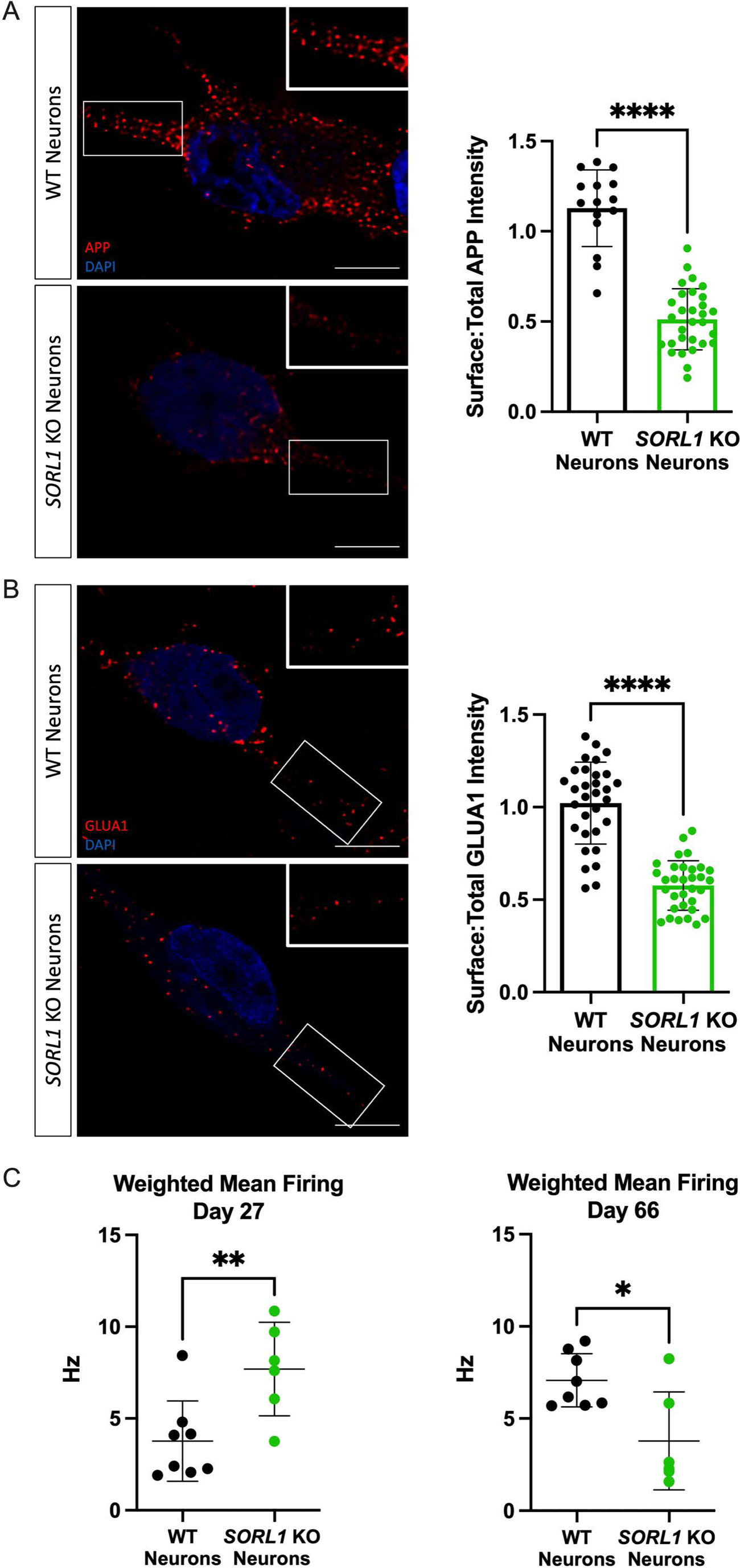
Loss of *SORL1* expression impairs recycling to the cell surface. **a-b)** *SORL1*KO neurons show reduced cell surface expression of APP **(a)** and GLUA1 **(b)**. Representative immunofluorescent images of WT and *SORL1*KO neurons labeled with antibodies for APP **(a)** (red) and GLUA1 **(b)** (red). Scale bar: 5μm. Intensity of APP and GLUA1 measured using ImageJ software. Data is presented as a ratio of surface intensity to total intensity. 2 WT and 2 *SORL1*KO clones were used in these experiments. 16 images per clone per genotype were analyzed. Data represented as mean ± SD. Normally distributed data was analyzed using parametric two-tailed unpaired t test. Significance was defined as a value of *p < 0.05, **p < 0.01, ***p < 0.001, and ****p < 0.0001. **(c)** Multielectrode array (MEA) analysis of WT and *SORL1*KO neurons at early (d27) and late (d66) time points of differentiation. 1 WT and 1 *SORL1*KO clone was used for these experiments. Data represented as mean ± SD. Data was analyzed using parametric two-tailed unpaired t test. Significance was defined as a value of *p < 0.05, **p < 0.01, ***p < 0.001, and ****p < 0.0001.

### *SORL1* overexpression enhances endosomal recycling

Defects in cell surface recycling have severe consequences in neurons, especially as these processes are necessary for healthy neuronal function and enhancing pathways that promote endosomal recycling in neurons may be beneficial. Using our *SORL1*OE neurons we analyzed recycling function using the transferrin recycling assay and observed that *SORL1*OE neurons showed significantly faster transferrin recycling (Figure 6A). We next tested whether colocalization of cargo with recycling endosomes and cell surface recycling was altered between *SORL1*OE and WT neurons. Interestingly, the size of Rab11+ recycling endosomes was significantly smaller in *SORL1*OE neurons (Figure 6B) possibly indicating that increased *SORL1* expression is clearing cargo more rapidly from this compartment. We observed a significant increase in localization of cargo with Rab11+ recycling endosomes (Figure 6C-E). While this result was initially surprising, as we also saw increased colocalization with Rab11+recycling endosomes in our *SORL1*KO neurons (Figure 4C-E), we further documented a significant increase of APP and GLUA1 on the cell surface compared to WT neurons with only endogenous *SORL1* expression (Figure 6F and 6G), as opposed to decreased APP and GLUA1 localization on the cell surface in *SORL1*KO neurons (Figure 5A and 5B). These results suggest that cell surface trafficking via a Rab11 pathway is enhanced by increased *SORL1* expression and that a crucial action of SORLA is the trafficking out of recycling endosomes. Thus, our data support a critical role for SORLA for trafficking cargo from recycling endosomes to the cell surface. In addition, we show for the first time that SORLA levels may regulate cell surface recycling of AMPA receptor subunits in human neurons.

**Figure 6.**
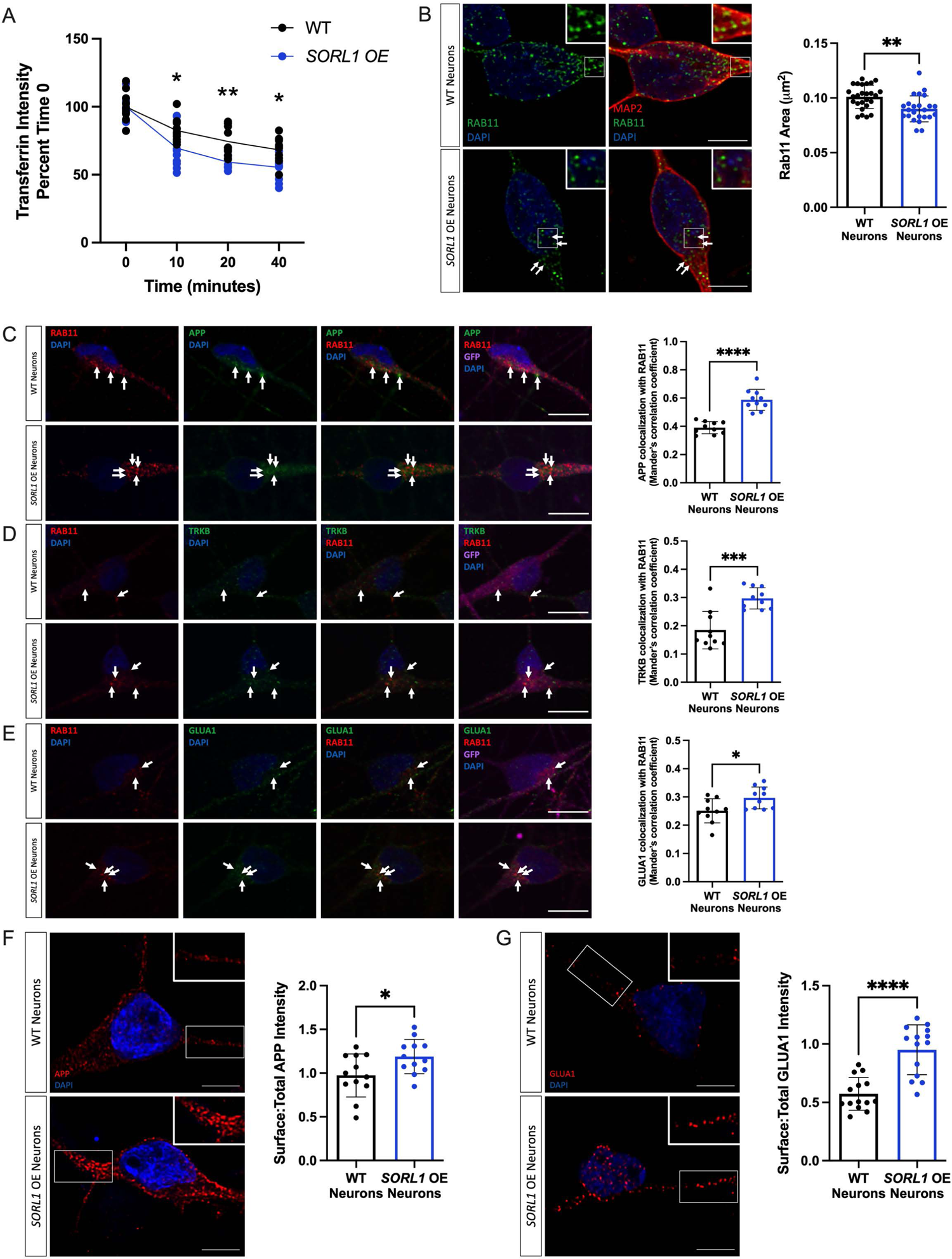
Overexpression of *SORL1* enhances endosomal recycling. **a)** *SORL1*OE neurons show faster rate of transferrin recycling. Quantification of fluorescence intensity of intracellular transferrin at different time points after treating cells with Alexa Fluor 647-conjugated transferrin for 15 mins, using ImageJ software. Data represented as percent of time 0 fluorescence intensity. 1 cell line of each genotype (WT vs. SORL1 OE) were used for these experiments. 10 images per genotype were analyzed. **(b)** *SORL1*OE neurons show reduced size of recycling endosomes. Representative immunofluorescent images of WT and *SORL1*OE neurons labeled with Rab11 (green) and MAP2 (red) showing smaller Rab11 positive recycling endosomes in *SORL1*OE neurons. Nuclei counterstained with DAPI (blue). Quantification of Rab11+ recycling endosome size performed using Cell Profiler software and represented as area of Rab11+ vesicles. Scale bar: 5μm. 1 cell line of each genotype (WT vs. SORL1 OE) were used for these experiments. 26 images per genotype were analyzed. Representative immunofluorescent images of WT and *SORL1*OE neurons showing increased colocalization of **(c)** APP (green), **(d)** TRKB (green) and **(e)** GLUA1 (green) with Rab11 (red) positive recycling endosomes. *SORL1*OE neurons and controls have endogenous GFP expression due to the piggybac vector system. GFP fluorescence is pseudo-colored (Far-red) and was used to outline cell bodies. Quantification of colocalization with Rab11 represented as Mander’s Correlation Co-efficient (MCC). Scale bar: 10μm. 1 cell line of each genotype (WT vs. SORL1 OE) were used for these experiments. 10 images per genotype were analyzed. Representative immunofluorescent images of WT and *SORL1*OE neurons showing increased cell surface expression of **(f)** APP (red) and **(g)** GLUA1 (red) in *SORL1*OE neurons. Scale bar: 5μm Fluorescence intensity of APP and GLUA1 measured using ImageJ software. Data is presented as a ratio of surface intensity to total intensity. Nuclei counterstained with DAPI. 1 cell line of each genotype (WT vs. SORL1 OE) were used for these experiments. 12-14 images per genotype were analyzed. Data represented as mean ± SD. Data was analyzed using parametric two-tailed unpaired t test and two-way ANOVA. Significance defined as a value of *p < 0.05, **p < 0.01, ***p < 0.001, and ****p < 0.0001.

### *SORL1* depletion affects gene expression

To determine a more global effect of chronic *SORL1* loss in human neurons, we performed bulk RNA sequencing of *SORL1*KO neurons compared to WT neurons. Interestingly, we observed that there were significantly more down-regulated genes in *SORL1*KO neurons than upregulated ones (Supplemental Figure 4B). While none of the cargo we explicitly studied in this work was differentially expressed, GO analysis showed that the top downregulated molecular function pathways in the *SORL1*KO cells were related to receptor-ligand activity and extracellular matrix organization (Figure 7A). The top upregulated molecular function pathways were related to ion channel activity (Figure 7B). To understand these data in the context of an integrated network, we used an analysis method that infers ligand receptor interactions from bulk RNA-seq data(Ramilowski et al., 2015; Wang et al., 2020). We observed several nodes of altered receptor-ligand interactions that indicate altered cell surface recycling and neurotrophic activity (Figure 7C). These include alterations in β-integrin signaling, which is consistent with previous work showing reduced β-integrin on the cell surface in *SORL1*KO cancer cells(Pietila et al., 2019), and altered interactions in ephrins/ephrin receptors, also corroborating previous work implicating *SORL1* expression in ephrin signaling and synapse regulation(Huang et al., 2017). Our analysis also showed nodes with alterations in nerve growth factor/nerve growth factor receptor (NGF/NGFR) and fibroblast growth factor/fibroblast growth factor receptor (FGF/FGFR) signaling, indicating alterations in neurotrophin and growth factor signaling and suggesting that the presence of endosomal traffic jams in *SORL1*KO neurons may ultimately impact multiple pathways important for neuronal health and development. Because we documented a decrease of GLUA1 puncta on the cell surface and observed altered neuronal activity on MEAs, we further mined our RNA-seq data for genes involved in synaptic function. Interestingly when we performed enrichment analysis, we observed that pathways associated with synaptic function were upregulated in *SORL1*KO neurons (Supplemental Figure 5).

**Figure 7.**
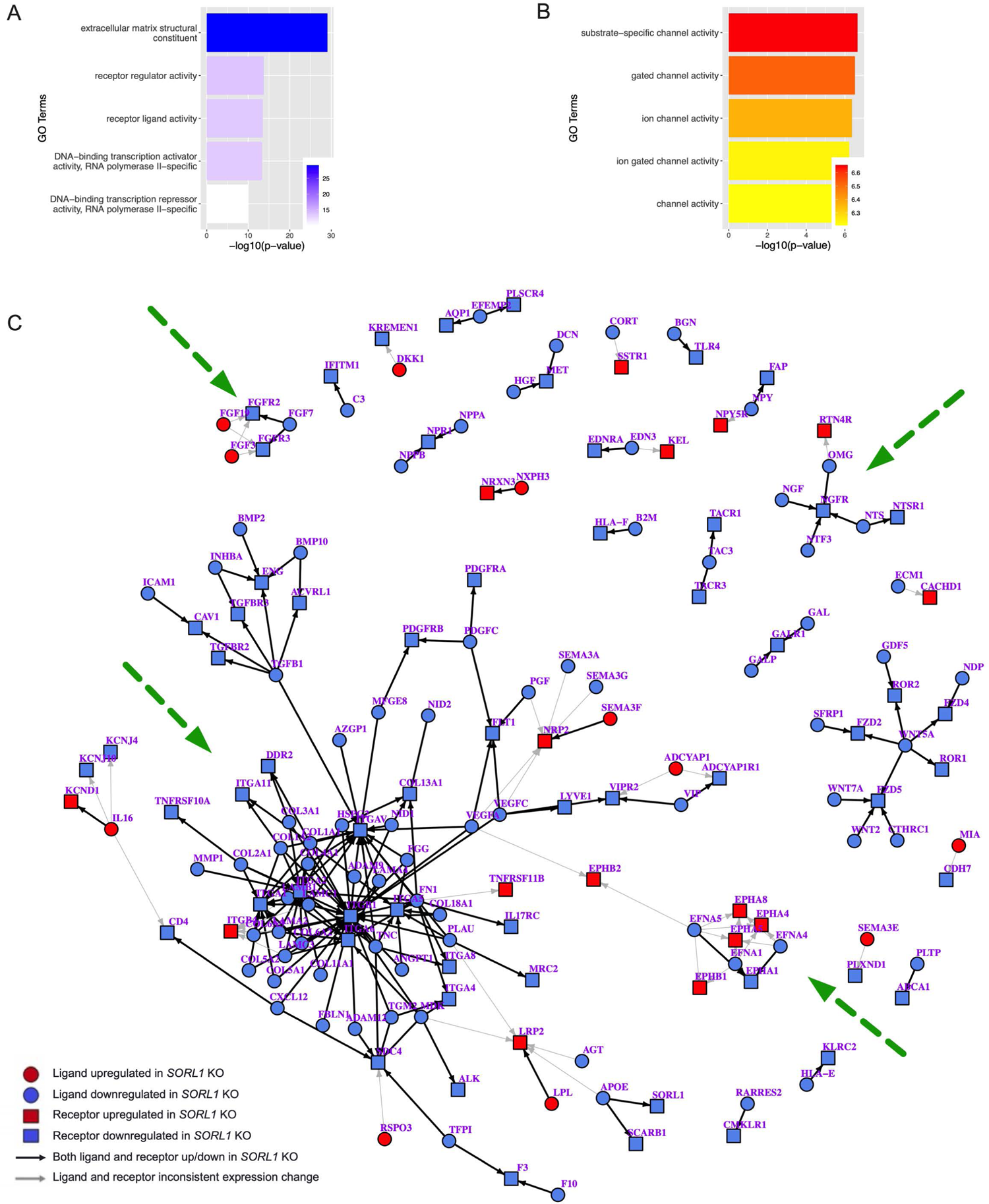
Analysis of bulk RNA-sequencing data indicates alterations in cell surface and extracellular trafficking, receptor-ligand and channel activity. Gene ontology analysis of DEGs in WT and *SORL1*KO neurons. Shown here are the top upregulated **(a)** and downregulated **(b)** molecular function terms in *SORL1*KO neurons. GO annotation terms are listed on the y-axis, adjusted p-value is shown on the x-axis. **(c)** Ligand-receptor network changes in *SORL1*KO neurons, identified if genes are more than 1.5-fold increased or decreased in *SORL1*KO neurons with an adjusted p-value less than 0.05. Circles denote ligands, squares denote receptors, blue indicates genes expressed significantly lower in *SORL1*KO neurons, red indicates genes expressed significantly higher in *SORL1*KO neurons. Arrows point from ligand to receptor, denoting receptor-ligand interactions. Black arrows denote consistent expression changes between ligand and receptor, indicating that both genes in the pair are either upregulated or downregulated. Gray arrows denote inconsistent changes between ligand and receptor. Clusters impacted by cell surface recycling (β-integrins, ephrins) or neurotrophic signaling (FGF/FGFR, NGF/NGFR) are indicated by green dotted arrows. RNA was collected from 3 separate differentiations including a combination of two WT clones and two *SORL1*KO clones. Each sample includes 2-3 technical replicates.

## DISCUSSION

Trafficking through the endo-lysosomal network regulates intra-cellular location of proteins, dictating their homeostasis, function and influence on cellular physiology. Functional studies by our group and others document endosomal abnormalities in hiPSC-derived neuronal models of AD(Hung et al., 2021; Knupp et al., 2020; Kwart et al., 2019). Emerging from this evidence is the role of *SORL1* as an endosomal gene that plays essential roles in mediating cargo trafficking. Recent work has implicated *SORL1* as AD’s fourth causal gene(Olav M. Andersen, 2021; Scheltens et al., 2021), and of these genes it is the only one linked to the common late-onset form of the disease. Understanding *SORL1*’s function is paramount for understanding AD’s pathogenic mechanisms and for potential therapeutic interventions.

Acting as an adaptor molecule of the retromer trafficking complex, SORLA has already been pathogenically linked to AD by its role in recycling APP out of endosomes(Andersen et al., 2005; Herskowitz et al., 2012; Offe et al., 2006). This current work and our previous study(Knupp et al., 2020) shows that *SORL1* depletion leads to increased APP localization in early and recycling endosomes. By lengthening the residence time of APP in these endosomal compartments, accelerated amyloidogenic cleavage of APP occurs due to the close proximity of APP and BACE1(Sun and Roy, 2018). Indeed, loss of *SORL1* leads to the accumulation of Aβ peptides, an antecedent of ‘amyloid pathology’ (Andersen et al., 2005; Knupp et al., 2020; Rogaeva et al., 2007b). We hypothesized that loss of *SORL1* in neurons would impact other cargo important for healthy neuronal function. To test this hypothesis, in addition to APP, we examined localization of the neurotrophin receptor TRKB and the GLUA1 subunit of the AMPA receptor and also observed that these proteins are increased in early endosomes (Figure. 1). These cargo link to another key pathology of AD: neurodegeneration, a slowly progressive process that begins with synaptic dysfunction characterized by glutamate receptor loss, which then progresses to synaptic loss before ultimately, over years, leading to widespread neuronal cell death (Selkoe, 2002).

The early endosome is considered the central station in the sorting and trafficking of cargo throughout the many stations of the endo-lysosomal system. While the early endosome is the station that is affected first and foremost in AD, it is not surprising that a primary dysfunction in this central station will secondarily influence trafficking throughout the system including the recycling and degradative pathways. Indeed, SORLA was shown to traffic Aβ to lysosomes in neuroblastoma cells, a function that is impaired by an AD-associated variant(Caglayan et al., 2014). Our work presented in this study, along with other recent work(Hung et al., 2021) also supports a role for *SORL1* in lysosomal trafficking in neurons. We observe a decrease in the pH-sensitive fluorogenic substrate DQ RED BSA in our *SORL1*KO neurons and decreased localization of APP, TRKB, and GLUA1 in late endosomes and lysosomes in *SORL1*KO neurons and that this is reversed in *SORL1*OE neurons (Figure. 2, Figure. 3). We interpret our functional and colocalization data to suggest that loss of *SORL1* expression mainly affects trafficking of cargo to lysosomes, but our data does not rule out a role of SORLA in neuronal lysosome function. Hung et al., reported decreased Cathepsin-D activity in *SORL1* deficient neurons, suggesting that *SORL1* loss directly impacts lysosome function(Hung et al., 2021). While we did not observe a difference in Cathepsin-D localization to lysosomes, it is important to note that that LAMP1 only partially colocalizes with Cathepsin-D in neurons (Cheng et al., 2018). Furthermore, the loss of proteolytic activity evidenced by decreased intensity of DQ Red BSA in *SORL1*KO neurons may not be completely due to reduced trafficking to lysosomes but could be a result of abnormal lysosomal function as DQ Red BSA is internalized by a process called macropinocytosis wherein macropinosomes can be directly trafficked to lysosomes (Hamasaki et al., 2004; Lorenzen et al., 2010; Racoosin and Swanson, 1993). Interestingly, while we did not observe an effect on DQ Red BSA at a shorter time point in *SORL1*OE neurons, we did see increased fluorescence of this reagent at a 24-hour time-point (Figure. 3), suggesting that overtime, increased SORL1 expression impacts lysosome trafficking and/or function in cortical neurons.

Our experiments also point to *SORL1*’s role in cell surface recycling. By using a prototypical cargo, transferrin, we demonstrate a reciprocal role between loss and enhancement of *SORL1* expression in cell surface recycling. Specifically, we show that *SORL1*KO neurons have defects in transferrin recycling at both early (10-minute) and late (40-minute) time points while *SORL1*OE neurons have faster recycling at these time points (Figure 4, Figure 6). This suggests that SORLA functions in both fast and slow endosomal recycling. Our data further implicates the recycling pathway by showing that modulation of SORLA expression affects recycling endosome size and the amount of cargo (APP, TRKB and GLUA1) localized to recycling endosomes (Figure. 4, Figure. 6).

Endocytic recycling comprises returning cargo, primarily membrane proteins, to the cell surface(Cullen and Steinberg, 2018). We studied a canonical SORLA cargo, APP, and show that loss of *SORL1* expression results in reduced cell surface APP while enhanced expression increases cell surface APP (Figure 5). These results corroborate previous work showing that *SORL1* and SNX27 work to return APP to the cell surface(Das et al., 2016; Huang et al., 2016) in a human model.

We also show that *SORL1* plays a role in recycling glutamate receptors (Figure. 5) This finding is critically important as recent work indicates that in mouse cortical neurons, SORLA interacts with a neuronal specific retromer subunit, VPS26b, to promote recycling of glutamate receptors(Simoes S., 2021). In our cortical neurons when *SORL1* is depleted, there is a reduction of GLUA1 subunits on the cell surface and this may result in synaptic impairment. MEA data comparing *SORL1*KO and isogenic WT neurons shows alterations in weighted mean firing rate as neurons mature (Figure 5). Interestingly, we observed an increase in neuronal firing at an early time point and a significant decrease in firing at a later time point. *SORL1*KO mice live to adulthood but have been described to have some deficits in learning and memory that may also be age dependent(Glerup et al., 2013; Hojland et al., 2018). Some of these alterations could be explained by compensatory expression changes ion channels or synaptic genes induced by chronic loss of *SORL1* during the course of neuronal differentiation from pluripotent stem cells. Our RNA-seq data does show up-regulation of ion channels and channel activity (Figure. 7). Interestingly, when we further interrogated our RNA-seq data for pathways enriched in synaptic function using the SynGo database (Koopmans 2019), we observed an upregulation of differentially expressed genes in synaptic pathways (Supplemental Fig. 5). This data suggests that *SORL1*KO neurons may attempt to compensate for altered trafficking of synaptic receptors by upregulating gene expression.

Our unbiased transcriptomic screen further supported that neurotrophic signaling and cell surface recycling pathways are impacted by *SORL1* deficiency (Figure 7). The SORLA cytoplasmic tail has been shown to translocate to the nucleus and activate transcription in a reporter gene assay(Bohm et al., 2006). Despite this, distinct genes regulated by SORLA are not known. Rather than looking for a direct effect on gene regulation, our goal for the analysis was to determine the global effect of *SORL1* loss or overexpression on neuronal networks.

Indeed, our data does not show that the specific cargo proteins described here are differentially expressed. However, the analysis does indicate that loss of *SORL1* in human neurons impacts cell surface networks, including receptor ligand interactions in neurotrophic and growth factor pathways, β-integrin signaling, and ephrin signaling. This corroborates previous work and the altered networks we observe impact neuronal health, axonal guidance, and synapse formation(Huang et al., 2017; Huang et al., 2006; Pietila et al., 2019).

Importantly, enhancing SORL1 expression improves cell surface trafficking of GLUA1 (Figure 6). Trafficking of glutamate receptors is an event that is critical for preventing synaptic dysfunction and synaptic loss, thus our results link *SORL1* to AD’s early-stage neurodegenerative process. Since retromer-dependent glutamate receptor recycling has been shown to occur independent of APP(Temkin et al., 2017), our previous and current results suggest that *SORL1* mutations can, at least in principle, drive two key AD pathologies, amyloid pathology and synaptic pathology, through parallel mechanisms(Small and Petsko, 2020).

We summarize our findings in Figure. 8. However, our study has certain limitations. For example, our results encompass only one human genome. Future studies will benefit from looking at *SORL1* deficiency or overexpression in multiple human genetic backgrounds. Furthermore, in this work we are describing purely neuronal phenotypes although *SORL1* is expressed in other CNS cells. Future work looking at cell-autonomous and non-cell autonomous mechanisms of *SORL1* depletion or overexpression in human glial or brain organoid models will be informative.

**Figure 8.**
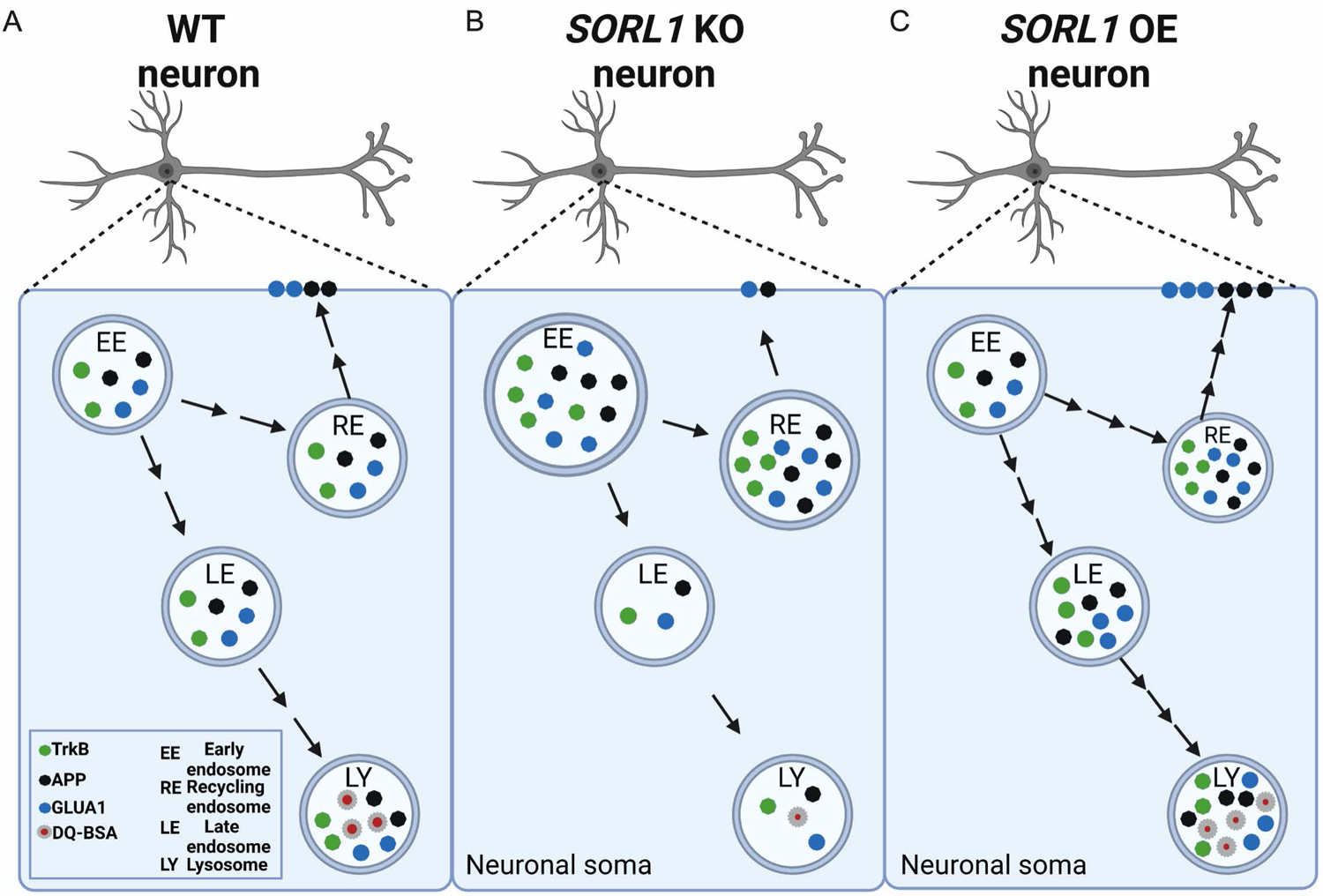
Model of modulation of SORL1 expression in hiPSC-derived cortical neurons. Modulation of SORLA expression regulates endosomal recycling and degradative pathways in hiPSC-derived neurons: The early endosome is a sorting hub where various cargo can be trafficked to degradative or cell surface recycling pathways in neurons (A). In this study, we mapped trafficking of three important neuronal cargo under conditions of depletion (B) or enhancement (C) of the AD risk gene SORL1/SORLA. As depicted in panel B, our data suggest that modulation of SORL1 expression significantly impacts the neuronal recycling pathway as we observe increased size and accumulation of cargo in recycling endosomes in SORL1 KO neurons compared to isogenic WT cells. Specifically, loss of SORL1 impacts recycling to the cell surface because, while there is increased cargo in recycling endosomes, there is a reduction of this cargo on the neuronal surface. Loss of SORL1 also impacts the degradative pathway out of the early endosome as there is a reduction of cargo in late endosome and lysosomes. As depicted in panel C, enhancement of SORL1 expression reciprocally impacts these pathways. SORL1 OE neurons also have an increase in cargo in recycling endosomes, but unlike KO cells, this results in more cell surface trafficking of neuronal cargo, leading to reduced stress (small size) of recycling endosomes. Additionally, SORL1 OE enhances trafficking out of the early endosome towards the degradative pathway as well. Taken together, our data solidifies the AD risk gene SORL1 as a key modulator of neuronal endosomal trafficking. Created with Biorender.com.

## CONCLUSIONS

In this work, we report that *SORL1* depletion affects endosomal trafficking by retaining cargo in early and recycling endosomes and impacts cell surface recycling and lysosomal trafficking of neuronal cargo. In particular, we demonstrate that *SORL1* expression in neurons affects cell surface localization of GLUA1, a phenotype that may ultimately impact synaptic dysfunction and neurodegeneration in AD. Interestingly, increasing *SORL1* expression enhances endosomal recycling and increases cell surface GLUA1. While the secondary downstream effects induced by *SORL1* depletion in the endo-lysosomal system are interesting and likely relevant to AD’s ultimate pathogenesis, from a therapeutic perspective it is best to target *SORL1*’s primary defect, which seems to localize to the endosomal recycling pathway. Interestingly, recent biomarker studies suggest that defects in retromer-dependent endosomal recycling occur in a majority of patients with ‘sporadic’ AD(Simoes et al., 2020), suggesting that the observed *SORL1*-induced defects may generalize across early and late onset forms of the disorder. Collectively, our results support the conclusion that *SORL1*, and the retromer-dependent pathway in which it functions, is a valid therapeutic target and interventions directed at this pathway may ameliorate endosomal recycling defects that seem to act as, at least, one primary driver of AD.

## ABBREVIATIONS

AD: Alzheimer’s Disease

SORL1: Sortilin-related receptor 1

SORLA: Sortilin-related receptor with A-type repeats

APP: Amyloid Precursor Protein

TRKB: Tropomyosin Related Kinase B

GLUA1: Glutamate receptor subunit AMPA1

AMPA: α-amino-3-hydroxy-5-methyl-4-isoxazolepropionic acid

OE: Overexpression

KO: Knock-out

WT: Wild-type

LAMP1: Lysosome-associated membrane glycoprotein 1

M6PR: mannose-6-phosphate receptor

GGA: Golgi-Localized g-Ear-Containing ARF-Binding

Rab: Ras-related protein.

## DECLARATIONS

### Ethics Approval and Consent to Participate

Not applicable

### Consent for Publication

Not applicable

### Availability of Data and Material

All raw and processed RNA-seq data has been deposited at the NCBI Gene Expression Omnibus (GSE180793). The data sets used and/or analyzed during the current study are available from the corresponding author on reasonable request.

### Availability of Supporting Material

All supporting data and material in the current study are available from the corresponding author on reasonable request.

## Acknowledgements

We thank Dr. Harald Frankowski, Dr. Yoshito Kinoshita, Ms. Shannon Rose, Ms. Eden Cruickshank and all members of the Young Laboratory for critical discussions and feedback during the preparation of this manuscript. We also thank Dr. Scott A. Small and Dr. Gregory A. Petsko for critical comments, discussions and feedback on this work. We would like to acknowledge the UW SLU Cell Analysis Facility and the Garvey Imaging Core at the UW Institute for Stem Cell and Regenerative Medicine. This work was supported by a NIH grant (R01AG062148) and a BrightFocus Foundation grant (A2018656S) to J.E.Y., a Biogen Sponsored Research Agreement to J.E.Y., a Retromer Therapeutic Sponsored Research Agreement to J.E.Y. and NIH traning grant (T32AG052354) to A.K. and a generous gift from the Ellison Foundation (to UW).

## Author Contributions

Conceptualization: J.E.Y., S.M, A.K. Microscopy analysis: S.M, A.K., and D.W.H. RNA-seq analysis and bioinformatics: Y.W and A.K. Methodology: S.M., A.K., M.S., C.A.W., C.K. Writing-Original Draft: S.M., A.K. and J.E.Y. Writing-Reviewing and editing: J.E.Y., S.M., A.K. Funding Acquisition: J.E.Y. and A.K. Supervision: J.E.Y. All authors read and approved the final manuscript.

## Supplementary figures

**Supplementary figure 1.**
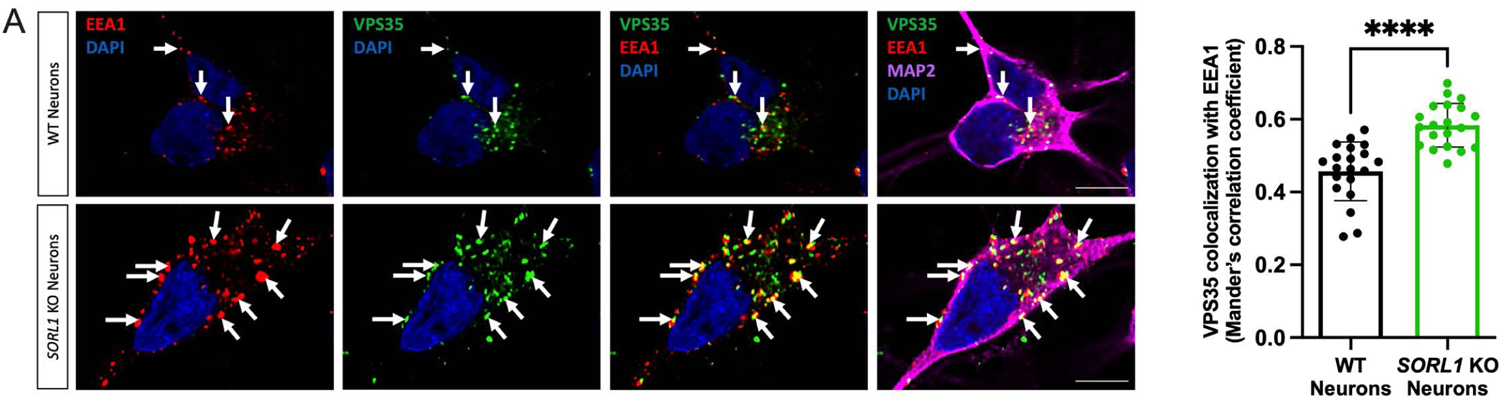
Loss of *SORL1* expression leads to increased VPS35 localization in early endosomes. **(a)** Representative immunofluorescent images of WT and *SORL1*KO neurons showing increased colocalization of VPS35 (green) with EEA1 (red). All neurons were immunolabeled with MAP2 (far-red) and counterstained with DAPI (blue). Scale bar: 10μm. In all cases, quantification of colocalization was represented as Mander’s correlation co-efficient (MCC). 1 WT and 2 SORL1KO isogenic clones were used for these experiments and 10 images per clone per genotype were analyzed. Data represented as mean ± SD. Significance was determined using parametric two-tailed unpaired t test and was defined as a value of *p < 0.05, **p < 0.01, ***p < 0.001, and ****p < 0.0001.

**Supplementary figure 2.**
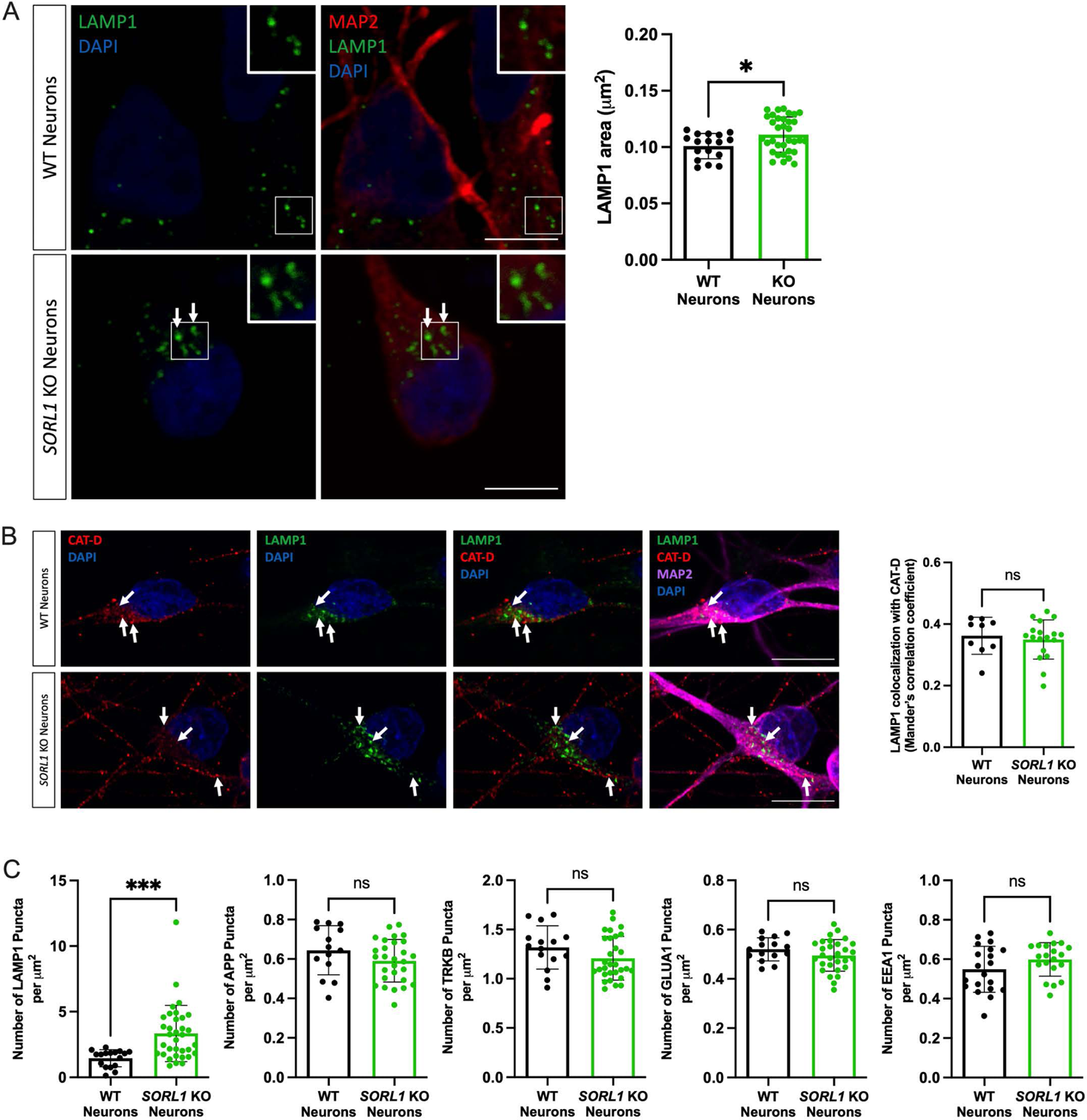
**a)** *SORL1*KO neurons show larger lysosome size and increased lysosome number. Representative immunofluorescent images of WT and *SORL1*KO neurons labeled with LAMP1 (green) and MAP2 (red) showing increased LAMP1 positive vesicle size in *SORL1*KO neurons. Quantification of LAMP1 size and number was performed using Cell Profiler software. LAMP1 size is represented as area of LAMP1 positive vesicles, and LAMP1 number is represented as number of LAMP1 positive vesicles per square micron of cell area. Scale bar: 5μm **(b)** *SORL1*KO neurons show no change in colocalization of lysosomes with the lysosomal enzyme Cathepsin D. Representative immunofluorescent images of WT and *SORL1*KO neurons labeled with antibodies for LAMP1 (green), Cathepsin D (red) and MAP2 (Far-red) showing no alteration in colocalization of Cathepsin-D with LAMP1 in *SORL1*KO neurons. Nuclei counterstained with DAPI (blue). Scale bar: 10μm Quantification of colocalization of LAMP1 with Cathepsin-D represented as Mander’s correlation coefficient (MCC). 10-20 images were analyzed per genotype. Two isogenic clones of each genotype were used in all experiments. Data represented as mean ± SD. Normally distributed data was analyzed using parametric two-tailed unpaired t test. Significance was defined as a value of *p < 0.05, **p < 0.01, ***p < 0.001, and ****p < 0.0001.

**Supplemental figure 3.**
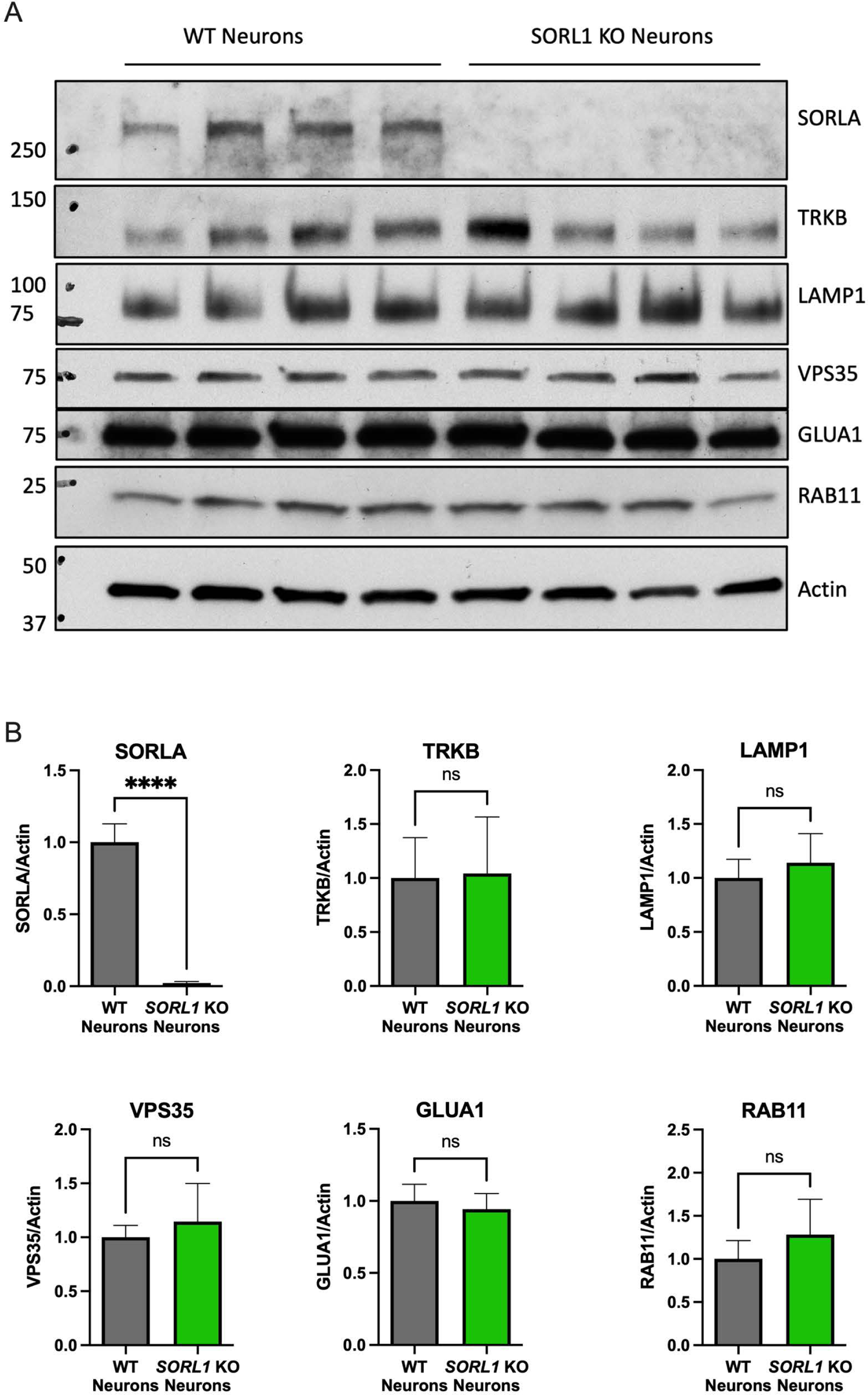
Loss of SORL1 does not change protein expression of the compartments or cargo analyzed in this study as analyzed by Western blot. Representative blots in (a), quantification in (b).

**Supplemental figure 4.**
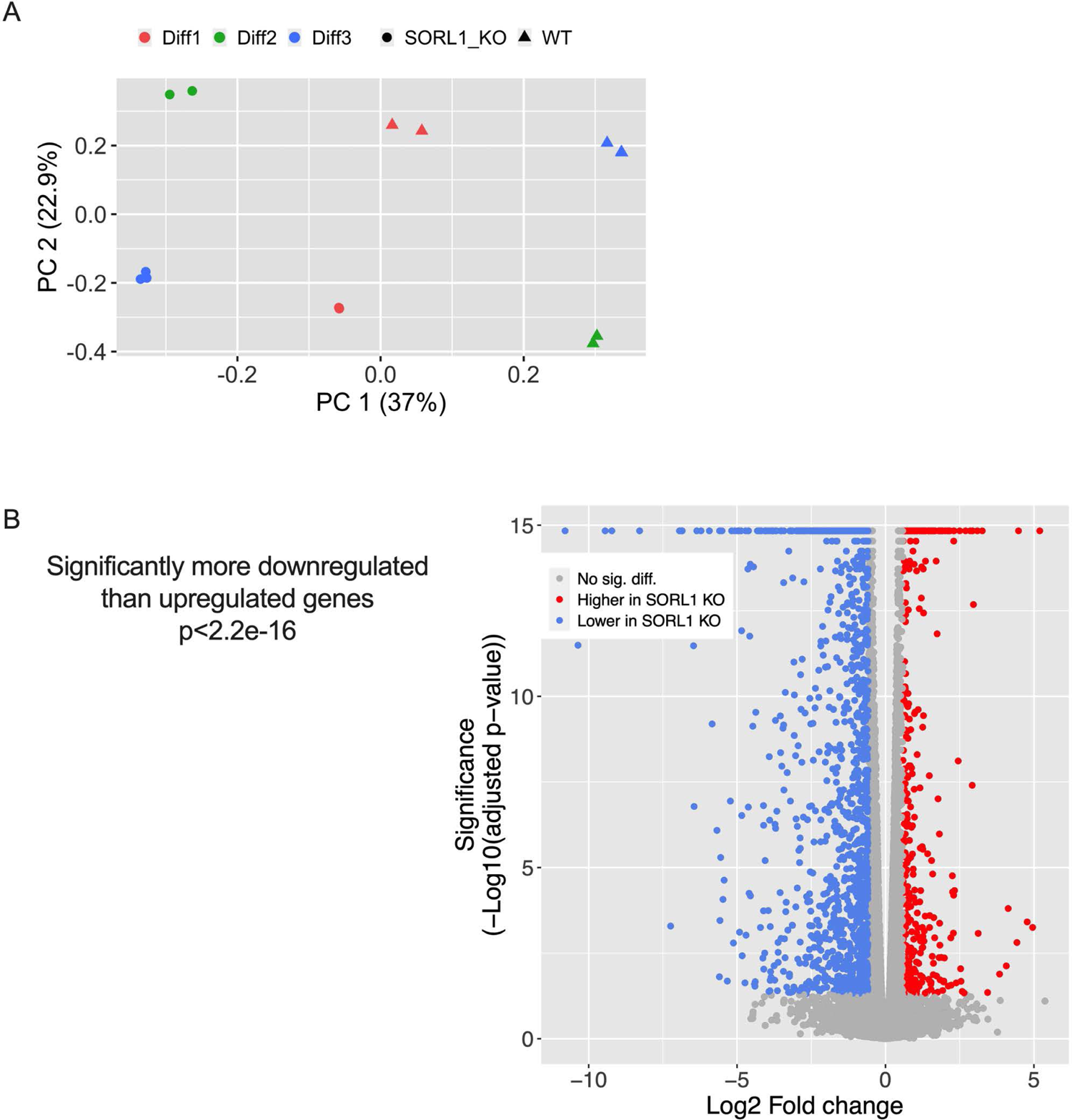
**a)** Principal component analysis (PCA) plot of all RNAseq samples using all expressed genes. Samples are color coded by differentiation batch. Triangles represent WT samples, circles represent *SORL1*KO. Genotype accounts for the highest variance (37%, PC1, x-axis). **b)** Volcano plot. Log2 fold change between *SORL1*KO and WT is shown along the x-axis. Statistical significance is shown along the y-axis, and is measured by adjusted p-value. Genes upregulated in *SORL1*KO neurons are shown by red circles, genes downregulated in *SORL1*KO neurons are shown by blue circles. We observed 6643 DEGs, with 2819 upregulated and 3824 downregulated. There are significantly more down regulated genes than upregulated genes (p<2.2e-16). Grey circles represent genes that are not significantly differentially expressed.

**Supplemental figure 5.**
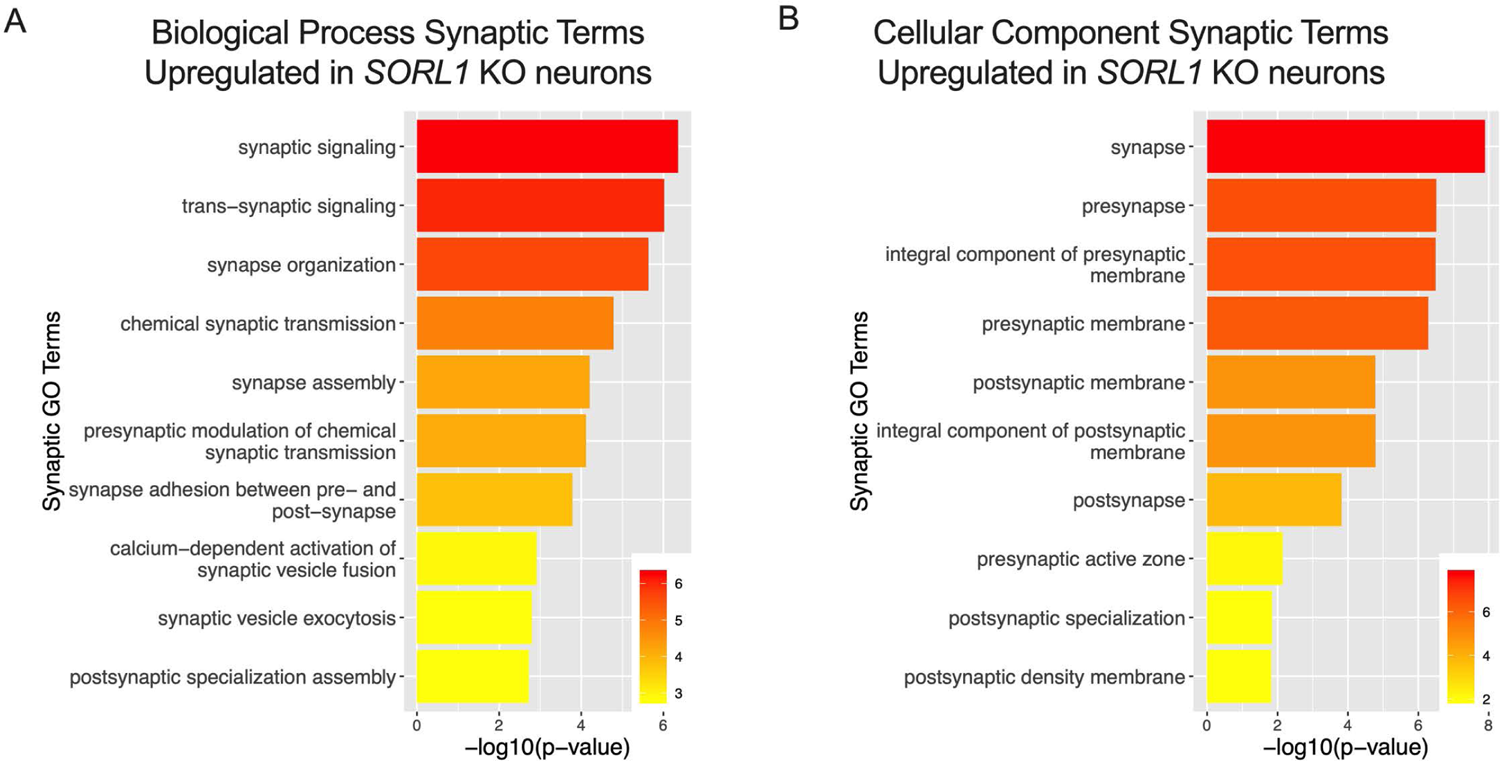
Loss of *SORL1* expression alters synaptic pathways. Analysis of bulk RNA-sequencing data indicates alterations in synaptic pathway functioning. We conducted gene ontology analysis of DEGs in WT and *SORL1*KO neurons using the SynGO synaptic annotation database. Shown here are the top upregulated and biological process and cellular component terms in *SORL1*KO neurons. GO annotation terms are listed on the y-axis, adjusted p-value is shown on the x-axis. No downregulated pathways in the SynGO database were shown to be enriched in *SORL1*KO neurons.

---

**Supplemental Table 1.** Summary of statistical analyses for data presented in this manuscript. In this table we present the statistical data that correspond to the experiments presented in the figures. This includes the group means, the difference between the means ± SEM, and the 95% confidence interval.

| Figure Number | Experimental Measure | Group A | Group B | Group A Mean | Group B Mean | Difference between means (B - A) ± SEM | 95% Confidence Interval |
| --- | --- | --- | --- | --- | --- | --- | --- |
| 1A | TRKB colocalization with EEA1 | WT Neurons | SORL1KO Neurons | 0.0799 | 0.1228 | 0.04290 ± 0.01042 | 0.02155 to 0.06425 |
| 1B | GLUA1 colocalization with EEA1 | WT Neurons | SORL1KO Neurons | 0.0496 | 0.1727 | 0.1231 ± 0.02257 | 0.07682 to 0.1693 |
| 2A | 6hr DQ Red BSA Intensity | WT Neurons | SORL1KO Neurons | 52.1 | 29.48 | -22.62 ± 2.684 | -28.09 to -17.15 |
| 2A | 24hr DQ Red BSA Intensity | WT Neurons | SORL1KO Neurons | 62.54 | 43.35 | -19.19 ± 3.608 | -26.58 to -11.80 |
| 2B | TRKB colocalization with RAB7 | WT Neurons | SORL1KO Neurons | 0.3102 | 0.2149 | -0.09530 ± 0.01234 | -0.1212 to -0.06937 |
| 2C | GLUA1 colocalization with RAB7 | WT Neurons | SORL1KO Neurons | 0.3828 | 0.286 | -0.09685 ± 0.02779 | -0.1538 to -0.03992 |
| 2D | APP colocalization with LAMP1 | WT Neurons | SORL1KO Neurons | 0.2983 | 0.1799 | -0.1184 ± 0.01951 | -0.1584 to -0.07844 |
| 2E | TRKB colocalization with LAMP1 | WT Neurons | SORL1KO Neurons | 0.071 | 0.04461 | -0.02639 ± 0.004876 | -0.03643 to -0.01635 |
| 2F | GLUA1 colocalization with LAMP1 | WT Neurons | SORL1KO Neurons | 0.1967 | 0.1588 | -0.03790 ± 0.02085 | -0.08060 to 0.004801 |
| 3A | 6hr DQ Red BSA Intensity | WT Neurons | SORL1OE Neurons | 30.31 | 27.84 | -2.473 ± 2.059 | -6.799 to 1.854 |
| 3A | 24hr DQ Red BSA Intensity | WT Neurons | SORL1OE Neurons | 32.61 | 39.73 | 7.123 ± 1.887 | 3.157 to 11.09 |
| 3B | APP colocalization with RAB7 | WT Neurons | SORL1OE Neurons | 0.7041 | 0.8008 | 0.09670 ± 0.02520 | 0.04375 to 0.1496 |
| 3C | TRKB colocalization with RAB7 | WT Neurons | SORL1OE Neurons | 0.44 | 0.5261 | 0.08610 ± 0.02417 | 0.03532 to 0.1369 |
| 3D | GLUA1 colocalization with RAB7 | WT Neurons | SORL1OE Neurons | 0.4287 | 0.5512 | 0.1225 ± 0.02485 | 0.07029 to 0.1747 |
| 3E | APP<br>colocalization<br>with LAMP1 | WT<br>Neurons | SORL1OE<br>Neurons | 0.1397 | 0.2949 | 0.1552 ±<br>0.02066 | 0.1118 to<br>0.1986 |
| 3F | TRKB<br>colocalization<br>with LAMP1 | WT<br>Neurons | SORL1OE<br>Neurons | 0.1539 | 0.2674 | 0.1135 ±<br>0.01501 | 0.08196 to<br>0.1450 |
| 3G | GLUA1<br>colocalization<br>with LAMP1 | WT<br>Neurons | SORL1OE<br>Neurons | 0.1353 | 0.3206 | 0.1853 ±<br>0.02279 | 0.1374 to<br>0.2332 |
| 4A - T0 | Transferrin<br>Intensity T0 | WT<br>Neurons<br>- T0 | SORL1KO<br>Neurons -<br>T0 | 100 | 100 | -4.167E-08 | -4.634 to<br>4.634 |
| 4A - T10 | Transferrin<br>Intensity T10 | WT<br>Neurons<br>- T10 | SORL1KO<br>Neurons -<br>T10 | 64.14 | 82.82 | -18.68 | -24.35 to -<br>13.00 |
| 4A - T20 | Transferrin<br>Intensity T20 | WT<br>Neurons<br>- T20 | SORL1KO<br>Neurons -<br>T20 | 58.44 | 67.48 | -9.038 | -13.67 to -<br>4.404 |
| 4A - T40 | Transferrin<br>Intensity T40 | WT<br>Neurons<br>- T40 | SORL1KO<br>Neurons -<br>T40 | 56.41 | 68.96 | -12.55 | -17.19 to -<br>7.920 |
| 4B | Rab11 Area | WT<br>Neurons | SORL1KO<br>Neurons | 0.06625 | 0.073 | 0.006747 ±<br>0.002882 | 0.0009338<br>to 0.01256 |
| 4C | APP<br>colocalization<br>with RAB11 | WT<br>Neurons | SORL1KO<br>Neurons | 0.2689 | 0.3147 | 0.04580 ±<br>0.01442 | 0.01626 to<br>0.07534 |
| 4D | TRKB<br>colocalization<br>with RAB11 | WT<br>Neurons | SORL1KO<br>Neurons | 0.1071 | 0.2277 | 0.1206 ±<br>0.02924 | 0.06071 to<br>0.1805 |
| 4E | GLUA1<br>colocalization<br>with RAB11 | WT<br>Neurons | SORL1KO<br>Neurons | 0.2605 | 0.3065 | 0.04595 ±<br>0.01599 | 0.01320 to<br>0.07870 |
| 5A | Surface:Total<br>APP Intensity | WT<br>Neurons | SORL1KO<br>Neurons | 1.128 | 0.5124 | -0.6157 ±<br>0.05825 | -0.7331 to -<br>0.4982 |
| 5B | Surface:Total<br>GLUA1<br>Intensity | WT<br>Neurons | SORL1KO<br>Neurons | 1.022 | 0.5769 | -0.4451 ±<br>0.04588 | -0.5368 to -<br>0.3534 |
| 5C | D27 Weighted<br>Mean Firing | WT<br>Neurons | SORL1KO<br>Neurons | 3.768 | 7.696 | 3.928 ± 1.266 | 1.169 to<br>6.687 |
| 5C | D66 Weighted<br>Mean Firing | WT<br>Neurons | SORL1KO<br>Neurons | 7.074 | 3.784 | -3.289 ±<br>1.101 | -5.688 to -<br>0.8903 |
| 6A - T0 | Transferrin Intensity T0 | WT Neurons - T0 | SORL1OE Neurons - T0 | 100 | 100 | -0.0000002 | -11.47 to 11.47 |
| 6A - T10 | Transferrin Intensity T10 | WT Neurons - T10 | SORL1OE Neurons - T10 | 82.52 | 69.57 | 12.95 | 1.483 to 24.42 |
| 6A - T20 | Transferrin Intensity T20 | WT Neurons - T20 | SORL1OE Neurons - T20 | 74.47 | 59.35 | 15.12 | 3.653 to 26.59 |
| 6A - T40 | Transferrin Intensity T40 | WT Neurons - T40 | SORL1OE Neurons - T40 | 68.31 | 55.43 | 12.89 | 1.419 to 24.35 |
| 6B | Rab11 Area | WT Neurons | SORL1OE Neurons | 0.101 | 0.09005 | -0.01099 ± 0.003199 | -0.01742 to -0.004558 |
| 6C | APP colocalization with RAB11 | WT Neurons | SORL1OE Neurons | 0.3903 | 0.5878 | 0.1975 ± 0.02717 | 0.1404 to 0.2546 |
| 6D | TRKB colocalization with RAB11 | WT Neurons | SORL1OE Neurons | 0.1846 | 0.2969 | 0.1123 ± 0.02416 | 0.06155 to 0.1631 |
| 6E | GLUA1 colocalization with RAB11 | WT Neurons | SORL1OE Neurons | 0.2509 | 0.2965 | 0.04560 ± 0.01822 | 0.007315 to 0.08389 |
| 6F | Surface:Total APP Intensity | WT Neurons | SORL1OE Neurons | 0.9734 | 1.189 | 0.2155 ± 0.09083 | 0.02710 to 0.4038 |
| 6G | Surface:Total GLUA1 Intensity | WT Neurons | SORL1OE Neurons | 0.5727 | 0.9505 | 0.3778 ± 0.06831 | 0.2374 to 0.5182 |
| Supplementary 1A | VPS35 colocalization with EEA1 | WT Neurons | SORL1KO Neurons | 0.457 | 0.5838 | 0.1268 ± 0.02249 | 0.08129 to 0.1724 |
| Supplementary 2A | LAMP1 area | WT Neurons | SORL1KO Neurons | 0.1008 | 0.1112 | 0.01032 ± 0.004276 | 0.001730 to 0.01892 |
| Supplementary 2B | Cathepsin-D colocalization with LAMP1 | WT Neurons | SORL1KO Neurons | 0.3621 | 0.3499 | -0.01219 ± 0.02558 | -0.06488 to 0.04051 |
| Supplementary 2C | Number of LAMP1 puncta per $\mu\text{m}^2$ | WT Neurons | SORL1KO Neurons | 1.452 | 3.339 | 1.887 ± 0.5339 | 0.8145 to 2.960 |
| Supplementary 2C | Number of APP puncta per $\mu\text{m}^2$ | WT Neurons | SORL1KO Neurons | 0.6442 | 0.5906 | -0.05352 ± 0.03725 | -0.1288 to 0.02175 |
| Supplementary<br>2C | Number of<br>TRKB puncta<br>per $\mu\text{m}^2$ | WT<br>Neurons | SORL1KO<br>Neurons | 1.318 | 1.208 | -0.1100 ±<br>0.06986 | -0.2509 to<br>0.03084 |
| Supplementary<br>2C | Number of<br>GLUA1 puncta<br>per $\mu\text{m}^2$ | WT<br>Neurons | SORL1KO<br>Neurons | 0.5202 | 0.4951 | -0.02509 ±<br>0.01880 | -0.06303 to<br>0.01285 |
| Supplementary<br>2C | Number of<br>EEA1 puncta<br>per $\mu\text{m}^2$ | WT<br>Neurons | SORL1KO<br>Neurons | 0.5488 | 0.5986 | 0.04975 ±<br>0.03228 | -0.01560 to<br>0.1151 |
| Supplementary<br>3B | SORLA/Actin | WT<br>Neurons | SORL1KO<br>Neurons | 1 | 0.02221 | -0.9778 ±<br>0.06426 | -1.135 to -<br>0.8205 |
| Supplementary<br>3B | TRKB/Actin | WT<br>Neurons | SORL1KO<br>Neurons | 1 | 1.043 | 0.04315 ±<br>0.3219 | -0.7444 to<br>0.8307 |
| Supplementary<br>3B | LAMP1/Actin | WT<br>Neurons | SORL1KO<br>Neurons | 1 | 1.141 | 0.1413 ±<br>0.1599 | -0.2499 to<br>0.5324 |
| Supplementary<br>3B | VPS35/Actin | WT<br>Neurons | SORL1KO<br>Neurons | 1 | 1.146 | 0.1459 ±<br>0.1848 | -0.3064 to<br>0.5982 |
| Supplementary<br>3B | GLUA1/Actin | WT<br>Neurons | SORL1KO<br>Neurons | 1 | 0.9428 | -0.05717 ±<br>0.07946 | -0.2516 to<br>0.1373 |

